# Comprehensive analysis and genome-wide association studies of biomass, chlorophyll, seed and salinity tolerance related traits in rice highlight genetic hotspots for crop improvement

**DOI:** 10.1101/2020.12.24.424354

**Authors:** Md Nafis Ul Alam, G.M. Nurnabi Azad Jewel, Tomalika Azim, Zeba I. Seraj

## Abstract

Farmland is on the decline and worldwide food security is at risk. Rice is the staple of choice for over half the Earth’s people. To sustain current demands and ascertain a food secure future, substandard farmland affected by abiotic stresses must be utilized. For rapid crop improvement, a broader understanding of polygenic traits like stress tolerance and crop yield is indispensable. To this end, the hidden diversity of resilient and neglected wild varieties must be traced back to their genetic roots. In this study, we separately assayed 15 phenotypes in a panel of 176 diverse accessions predominantly comprised of local landraces from Bangladesh. We compiled high resolution sequence data for these accessions. We collectively studied the ties between the observed phenotypic differences and the examined additive genetic effects underlying these variations. We applied a sophisticated fixed effect model to associate phenotypes with genotypes on a genomic scale. Discovered QTLs were mapped to known genes. Candidate genes were sorted by tissue specific gene expression profiles and protein level consequence of existing polymorphisms. Our explorations yielded 17 QTLs related to various traits in multiple trait classes. 12 identified QTLs were equivalent to findings from previous studies. Integrative analysis assumes novel functionality for 21 candidate genes on multiple evidence levels. These findings will usher novel avenues for the bioengineering of high yielding crops of the future fortified with genetic defenses against abiotic stressors.

## 1 Introduction

10 million hectares of arable land is lost to urbanization every year [1]. An accelerated wave of climate change damages the quality of the remaining landmass [2]. About half of the world’s currently estimated farmland is affected by abiotic stresses, most notably, salinity [3, 4]. Among cereal crops, rice is the most sensitive to abiotic factors and yield losses [5] but has the largest contribution to global food production. Rice feeds more people across the world than any other crop [6], accounting for up to 80% of daily calorie intakes of half the world’s population [7]. It is surmised that global rice production has to double by the year 2050 to combat these burdens imposed on food security [8, 9]. The insufficiency in finite arable land has to be overcome by increases in crop productivity. This must involve the introgression and propagation of stress tolerance traits to enable farming in soil less ideal for present day elite varieties.

With the help of high-resolution genotyping technologies, scientists endeavor to elucidate the molecular signature of complex polygenic traits. Multitudes of small genetic effects and their interactions with the surroundings underlie the quantitative assertion of these phenotypes. This limits our ability to refine them through direct breeding or molecular approaches. Many statistical and genome-wide association based studies (GWAS) are carried out for salinity tolerance [10, 11], chlorophyll [12] and yield [13] related traits in rice independently. Researchers often point out that complex traits are interlinked through their evolutionary origins and must be studied in conjunction to reach conclusive outcomes [14]. No recent GWAS study explores abiotic stress tolerance in relation to developmental and agronomic phenotypes in rice. Identification of major-effect QTLs for yield, tolerance and developmental attributes will continue to add layers of sophistication to the current understanding of plant biology. Characterization of causal polymorphisms will facilitate the introgression of salient QTLs into elite varieties with the help of sophisticated molecular technologies such as CRISPR Cas9 genome editing.

Bangladesh is the largest delta on the planet, endowed with fertile land, meandering rivers and a pronounced history of growing rice. To accelerate the improvements in cereal crop development, we elaborately phenotyped 176 local rice varieties from Bangladesh for 15 quantitative traits relating to plant biomass, chlorophyll content, tissue ion content, visual salt damage and seed properties. We incorporated next-generation sequencing data with high genomic coverage to estimate additive genetic effects and correlate the observed phenotypic values with genomic predictions. We tested over 4 million high-quality SNP markers for association signals separately for each trait. We cross-referenced our identified QTLs with functional annotations, gene expression profiles and allelic substitution effects. The compiled assessment of multiple traits belonging to several trait classes accentuated the genetic landscape influencing their effects. For identified QTLs, integrative analysis of auxiliary data enabled us to unravel novel gene roles.

## 2 Methods

### 2.1 Data and source code

All data, source code and shell commands are documented on GitHub at https://www.github.com/DeadlineWasYesterday/Cat-does-plant. The complete computational pipeline was executed in an HP Z840 workstation running on a 16-core Intel Xeon processor with 256GB of RAM.

### 2.2 Plant material and growth conditions

The 176 rice accessions were ordered from the IRRI seed bank to ascertain their identity with the 3,000-rice genome project [15]. Seeds were multiplied at BRRI (Bangladesh Rice Research Institute) fields where the day/night temperature was 32°/28°C with an average humidity of 72%. Supplementary table 1 lists the accession codes and metadata with local names and subpopulation information for all 176 varieties. Only 3 varieties belong to the USDA core (IRGC 31727, IRGC 27555 and IRGC 58736) and none are listed in the USDA mini-core collection. 106 accessions are traditional landraces that are cultivated locally in Bangladesh. The remaining 70 accessions are varieties that had been developed by BRRI from local landraces for local agronomy. With the exception of three (IRGC 49375, IRGC 126002 and IRGC 124432), all varieties had been originally collected from Bangladesh by the International Rice Research Institute. The map of Bangladesh was drawn in Adobe Illustrator and labelled using the ggplot2 and ggrepel libraries in R.

The phenotype screening for salinity tolerance at the seedling stage took place in a nethouse enclosure at an average day/night temperature of 31°C/27°C and approximately 72% relative humidity. The screening for salinity tolerance at seedling stage was carried out following the methods described by Amin et al. 2012 [16]. Sprouted seeds were sown in netted Styrofoam and floated in 3×3 (3 for control and 3 for stress) replicated PVC trays containing 60L Yoshida solution [17]. The positions of the 176 accession seedlings were randomized in each of the 6 trays. The germinated seeds were allowed to grow for 14 days. Then, NaCl stress was applied gradually starting from 4dS/m up to 12dS/m by 2dS/m increments every 24 hours in 3 of the stress-experiment trays. The solution level of the 3 control trays were maintained with water. Phenotypic measurements were taken after 16 days of stress application, when 90% of the leaves of the sensitive control (IR29) were damaged.

### 2.3 Phenotyping

After about 1 month of germination and 16 days from the first salt stress exposure for the stress condition plants, seedlings were systematically phenotyped. Fresh weight and length measurements were taken for whole roots and shoots. For root and shoot ion content data, plants were externally washed and oven dried at 70°C. Ground-up dried roots and leaves were treated with 1N HCl for 48 hours before being assayed in a flame photometer (Sherwood model 410, Sherwood, UK). Ion measurements are denoted in millimolar concentration. Chlorophyll content was measured in fully expanded third leaves. Leaf extracts of 1cm^2^ size were dissolved in 80% acetone in the absence of light. Absorbance at 645nm and 663nm wavelengths were measured. bThe formulae for chlorophyll A and B estimation were adapted from Yoshida et al. 1976 [17] as follows:

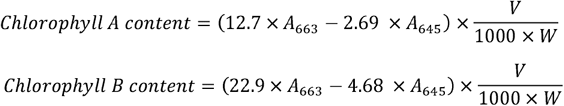

SES (Standard Evaluation System) scores were given to each plant on the basis of growth stunting, damage to leaves, leaf chlorosis, drying of biomass and plant vigor as described by Gregorio et al. 1997 [18, 19]. The data for seed traits were collected from seeds produced under normal conditions. Seed husks were removed by hand and dimensions of the kernels were measured using slide calipers. The average dimensions of 10 visually identical seeds from the same plant were registered as the working value for each replicate. Seed length is defined as the distance along the longest axis of the seed. In the axis perpendicular to seed length, seed width and height are defined as the shortest and longest distances respectively. The seed weight value is an alias for 1000 grain weight obtained by weighing 200 grains on a digital scale and scaling the value. All traits other than the SES score which is a factorial variable by definition, were logged as continuous numeric features.

### 2.4 Phenotype statistics

For the 10 traits measured in both conditions, heritability was measured by the formula 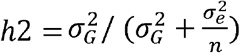 where 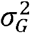 is the variance of the genotypes, 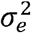 is the residual variance and *n* is the number of replicates. The variance components were calculated by including the genotype, treatment condition and interaction between genotype and condition as random effects in the *lmer* function from the lme4 package in R. Marker-based heritability was calculated using the kinship matrix K (details in section 2.7) in the *marker_h2* function from the heritability package in R. Broad-sense heritability was calculated by dividing the variance of the genotype means by the overall phenotypic variance. Correlation and GWAS studies involved phenotype means for three replicates. Linear correlation is expressed by the Pearson’s correlation coefficient. Genomic prediction for SNP effects on phenotype was calculated by ridge regression using the *mixed*.*solve* function from the rrBLUP package [20]. Assessment of data distributions was carried out by the SciPy module in Python. Phenotype transformation for GWAS was done by an ideal genotype derived warping function calculated by WarpedLMM [21].

### 2.5 Preprocessing genotype data

Next-generation sequencing data was compiled from the 3,000-rice genome project [15]. In the original study, libraries were created from young leaves and sequenced in the Illumina HiSeq2000 platform to generate 90bp paired-end reads. Raw sequence reads were aligned to the IRGSP 1.0 reference genome for *Oryza sativa sub. japonica* cultivar Nipponbare. The average sequencing depth was 14× with genome coverage and mapping rates over 90%. Publicly available VCF data from aligned reads were sourced for 176 varieties and compiled using VCFtools in the SAMtools suite. SNPs having phred-scaled quality scores below 30 were flagged as low-quality reads and considered missing data. Beagle 5.1 [22] was used for imputation of unphased genotypes. There were on average 2,930,739 missing genotypes out of 15,241,544 total marker calls per individual with a standard deviation of 492,558. For assessment of imputation accuracy, we prepared a test set having the original proportion of missing data by systematically removing high-quality reads from 1.85 million markers which had no missing reads. Unphased imputation accuracy was measured to be over 99.5% and the report can be found on the git repository. Working files were prepared using in-house python scripts. SNPs that originally had over 20% low quality reads were excluded from all studies. For GWAS, a minor allele frequency filter of 5% and 10% was applied to the whole population and the subpopulations respectively.

### 2.6 Population structure estimation

A combination of methods was employed for deciphering population structure. Principle components were calculated using R and clustered in two and three dimensions respectively using a k-means clustering algorithm. The first three principal components showed 38% explained variation in scree plots. 2D and 3D scatter plots were also plotted in R using the first three principal components and can be seen in the git repository. Bayesian maximum likelihood-based population structure estimation software fastStructure [23] implemented in python was used for assessing admixture between groups. K (number of subpopulations) values from 2 to 15 were tested and the *chooseK*.*py* function was used to select k = 3. Distances for the neighbor-joining tree were calculated in TASSEL 5 [24] and visualized by the web-based application iTOL (interactive Tree Of Life) [25].

### 2.7 Genome-wide association studies

The primary algorithm for association testing was BLINK [26] in the GAPIT software package [27]. The more popular and statistically robust compressed mixed linear model (CMLM) applied both in GAPIT and TASSEL 5 were also run to validate the findings (data in git repository).

BLINK employs two successive fixed effects models. The first model tests for associations and calculates p values and the second model iteratively improves the first model by exporting markers as covariates using Bayesian Information Criterion (BIC).

The first model is denoted:

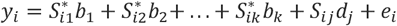

where *y*_*i*_ is the estimated observation for the *i*-th individual. 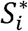 terms represent covariates named pseudo QTNs that start off as an empty set and subsequently are iteratively selected by the second model. terms are the corresponding effects for the pseudo QTNs. *S*_*ij*_is the genotype for the testing marker *j. d*_*j*_ is the effect of the *j*-th marker and *e*_*i*_is the residual error term having a distribution 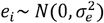.

The second model given below is similar but lacks the genotype term:

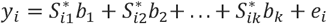

Markers are sorted by ascending order of p value and fitted in the second model one by one if considered significant after Bonferroni correction *and* found to not be in linkage disequilibrium (r^2^ > 0.7) with any marker already included in the equation. Model selection by BIC finds the best fit second model from all combinations and the covariates from that model are exported to the first model.

This complete process is repeated in BLINK until the second model no longer selects new covariates for inclusion in the first model and the first model is established as the final testing model.

To correct for population structure and cryptic relatedness, the kinship matrix K, calculated using GAPIT and the Q matrix having the first three principal components were fit in the first model as covariates. For the compressed mixed linear models fit for validation tests, the optimum compression level was selected and the same covariates were included. Q-Q plots and Manhattan plots were visualized by the qqman package in R [28].

An adjusted Bonferroni correction was applied to set the genome wide significant and suggestive thresholds. The formula used for setting the significance threshold was -log (1/*M*) where M is the total number of testing markers. In the whole population, the value was 6.63 for 4,283,120 markers that remained after the MAF filter, which was rounded down to 6.5 for use. In Indica and Aus subpopulations, there were 2,735,794 and 2,562,596 markers respectively and the significant threshold was set at 6.4 accordingly. For suggestive thresholds, effective number of markers were calculated, defined as those being in approximate linkage equilibrium across the genome. A pairwise LD cutoff of r^2^ = 0.2 with 50kb sliding windows and step size of 50kb in PLINK [29] was used to calculate the effective number of markers. The effective number was 384875, 255487 and 234071 for the whole, Indica and Aus sets respectively. The resulting suggestive threshold values were 5.37, 5.4 and 5.58, from which a common down-rounded value of 5.0 was used.

### 2.8 Gene expression and downstream analysis

Gene annotations were collected from the MSU7 [13] and RAP-DB [30] databases. RNA-seq expression data for 12 tissues and 4 seedlings were collected from MSU7. The sequence reads had been mapped by TopHat [31] and the RNA-seq libraries had been calculated with Cufflinks [32]. Genome-wide LD decay was plotted using PopLDdecay [33]. Heatmaps were generated using *pyplot* functions from the Matplotlib library in Python. For hierarchical clustering, the *hclust* and *heatmap* functions in R were used. An elaborate pipeline written in Python and R for the determination of SNP substitution effects is documented in GitHub. One-way ANOVA and t-tests were conducted using the SciPy library. The Benjamini-Hochberg FDR correction was conducted using the statsmodels library in Python.

## 3 Results

### 3.1 Population structure estimation

The panel of 176 accessions consists of varieties from the 3,000-rice genome project that had been sourced from Bangladesh. These varieties predominantly belong to the Indica and Aus subgroups. Bangladesh is the largest delta on Earth with bountiful rivers draining into the northeastern part of the Indian Ocean by way of the Bay of Bengal. Drainage of enormous amounts of sediments and erosion of upland rocks have promoted highly heterogeneous soil formation between floodplain land, intermittently wet plains, saline coasts, hill areas and uplifted terraces. In many places across the country where drought is common and along the southern coast where soil is saline, local landraces are predominantly farmed as they perform better than the sensitive elite varieties. Adaptation to uniquely materialized soil, locality-specific cultural farming practices and selection pressures from the environment embellish immense diversity to these genotypes. Figure 1 shows 114 known geolocations from supplementary table 1 where the labelled varieties have been historically cultivated.

**Figure 1.**
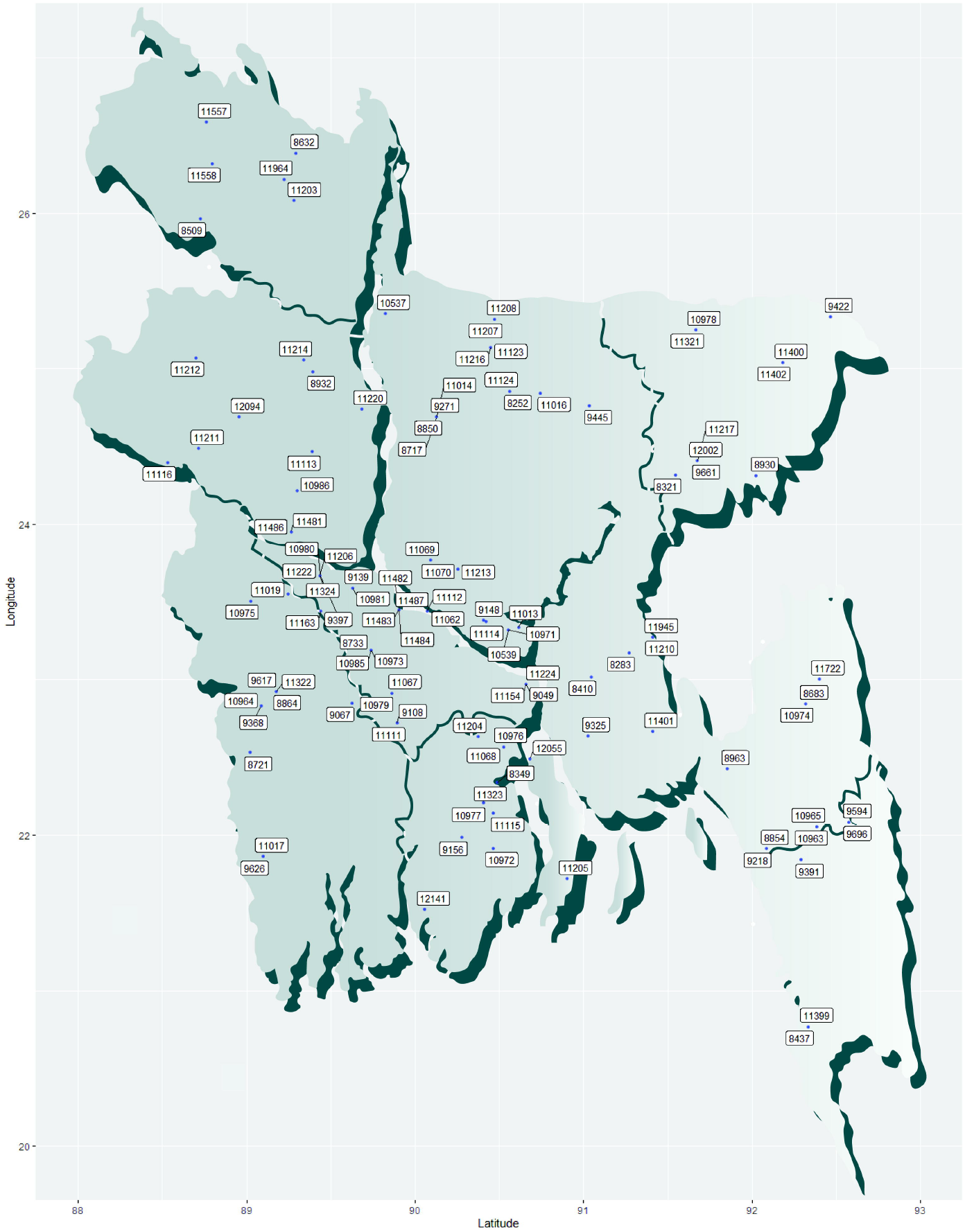
Map of Bangladesh showing the locations where 114 out of 176 rice varieties from this study have been traditionally propagated and cultivated. The labels in the image are the IRIS codes from supplementary table 1 lacking the common ‘IRIS_313-’ prefix.

For population structure estimation, we first constructed a Neighbor-Joining tree with 4,140,858 biallelic markers that remained after applying a minor allele frequency filter of 0.05. The Python implementation of the Bayesian clustering program STRUCTURE named fastStructure estimated the appropriate number of subpopulations (K) to be 3. Three distinct clusters could also be visualized in a scatter plot of the first and second principal components. An admixture plot for 3 subpopulations and the NJ tree for the 176 individuals can be seen in Figure 2 (a) and (b) respectively. The red individuals belong to the Aus subpopulation and the green shaded individuals are members of the Indica subpopulation. We analyzed 2D and 3D scatter plots for the first three principal components to infer population structure. The unsupervised k-means clustering algorithm was implemented on the PCA data in two and three dimensions separately. The clustering results can be seen in Figure 2 (c) and (d). The subpopulations assigned to the individuals by fastStructure, the groupings calculated by the k-means algorithm, and that inferred by manual inspection were found to be identical. Population structure inferences from all sources have been tabulated in supplementary table 1. In the NJ tree and the PCA scatter plots, the Aus subpopulation was found to cluster more tightly than the individuals belonging to the Indica subgroup. The third subpopulation lodges only a handful of individuals that are grouped as aromatic rice varieties and are locally known as Basmati or Sadri rice. Because of having a very small number of individuals, the aromatic subpopulation was excluded from subpopulation specific association testing along with three accessions that did not cluster with either subpopulation (Supplementary table 1).

**Figure 2.**
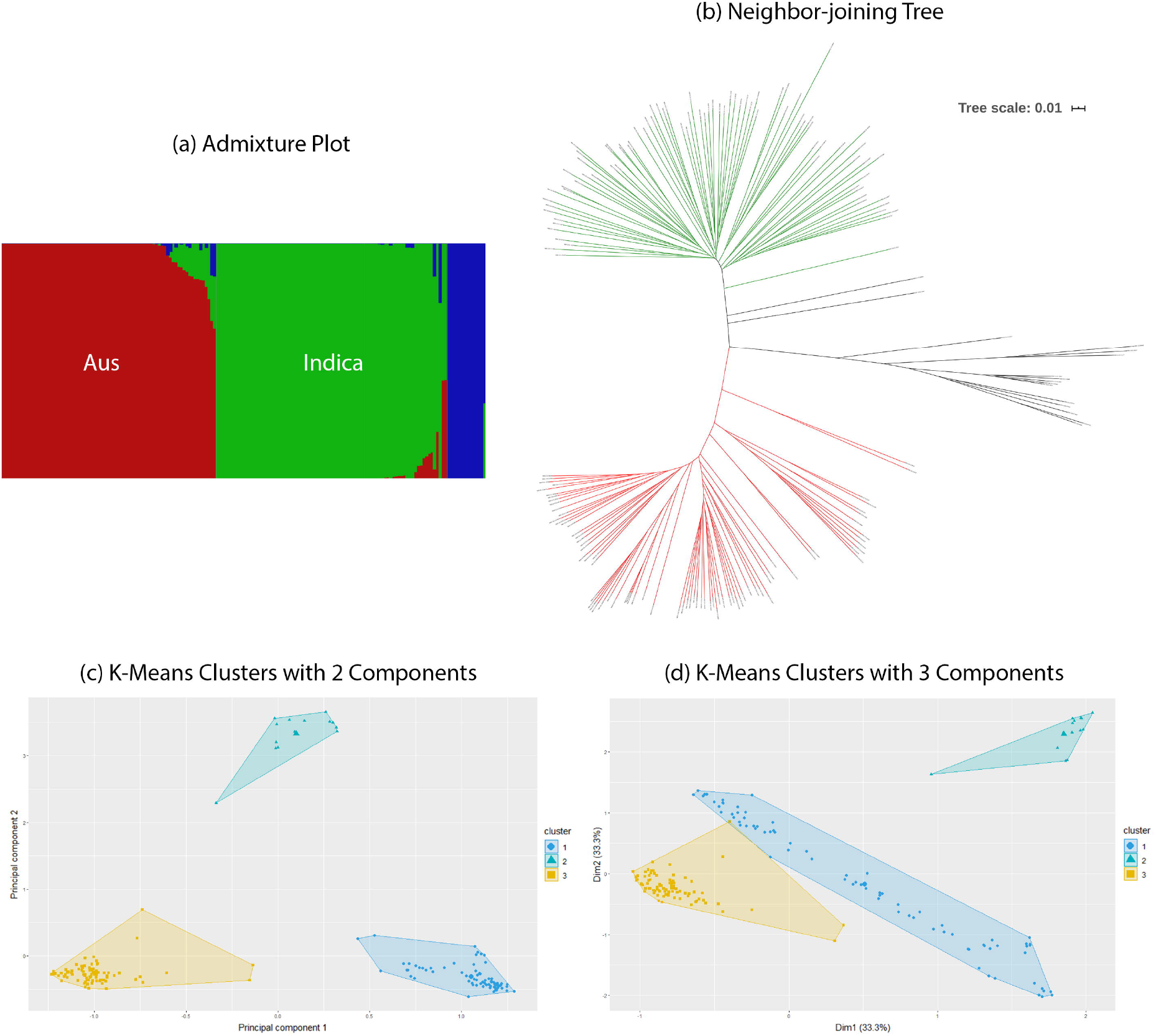
Population structure estimation results for 176 varieties. Three distinct subpopulations can be inferred from these diagrams. (a) An admixture plot calculated from fastStructure. (b) Neighbor-joining tree drawn from genotype data in TASSEL 5. The green individuals belong to the Aus subgroup, red individuals are from the Indica subgroup and black lines denote plants distant from these groups. (c) K-means clustering results of the first two principal components. (d) K-means clustering results of a three-dimensional plot of the first three principal components.

### 3.2 Comprehensive analysis of phenotypes

We have catalogued 14 basic traits that we grossly assign to 4 categories: plant biomass, tissue ion content, leaf chlorophyll content and seed properties. The 10 basic traits belonging to biomass, ion content and chlorophyll content categories were measured in both control and salt stress conditions. The seed traits were measured from plants grown under normal conditions. The 15th basic trait is the visual SES score for evaluating salt damage which by definition is only measured in stress condition. Simple mathematical operations on the plant biomass, leaf chlorophyll and seed related phenotypes gave rise to 14 additional derived phenotypes. Table 1 lists all basic and derived phenotypes with the formulations for each derivation from the basic traits. For the basic phenotypes measured in two treatment conditions, heritability was calculated using the variance component of the genotype and the residual variance. Variance components were obtained by estimating random intercepts for genotype groups, treatment conditions and treatment conditions within genotype groups. Heritability for the remaining basic phenotypes and all of the derived phenotypes were calculated using the genomic kinship matrix K. Marker derived heritability for all phenotypes and broad-sense heritability for applicable phenotypes can be found in supplementary table 2. The means of phenotype values recorded for every genotype is stored in supplementary table 3. Narrow-sense heritability could not be calculated from additive marker effects because the grand mean of the phenotypes could explain more variance than the *mixed*.*solve* prediction model. The linear Pearson’s correlation coefficients between the observed phenotypes are shown in the bottom diagonal of the heatmap in figure 3 and the correlation between the genotype effects for each phenotype calculated through genomic prediction occupies the upper diagonal of the heatmap. The correlation between the observed phenotypes and additive SNP effects are seen along the white diagonal of the map. All phenotype estimations made by genomic prediction can be found in supplementary table 4. An extended figure showing correlation coefficients between observed and estimated values of all 39 basic and derived phenotypes can be viewed in supplementary figure 1. A matrix of p values for the correlations shown in supplementary figure 1 is provided in supplementary table 5.

**Table 1:**
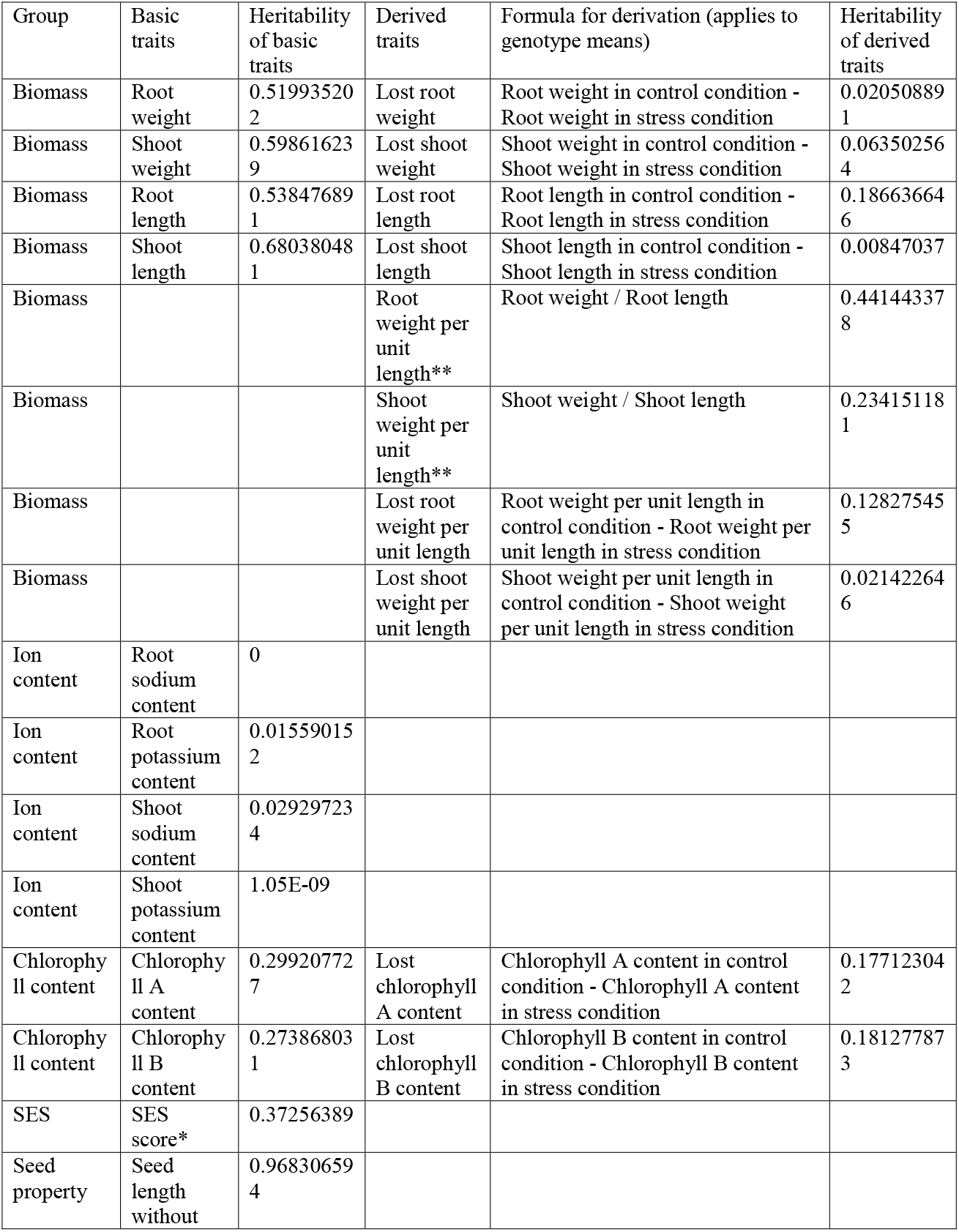
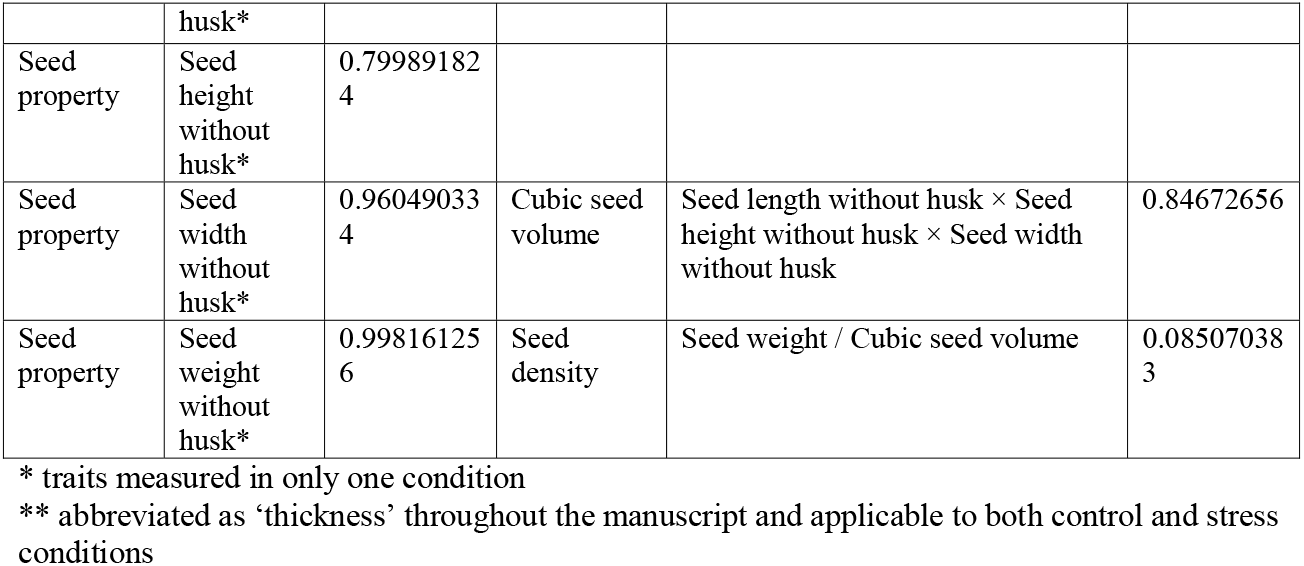
Basic and derived phenotypes.

**Figure 3.**
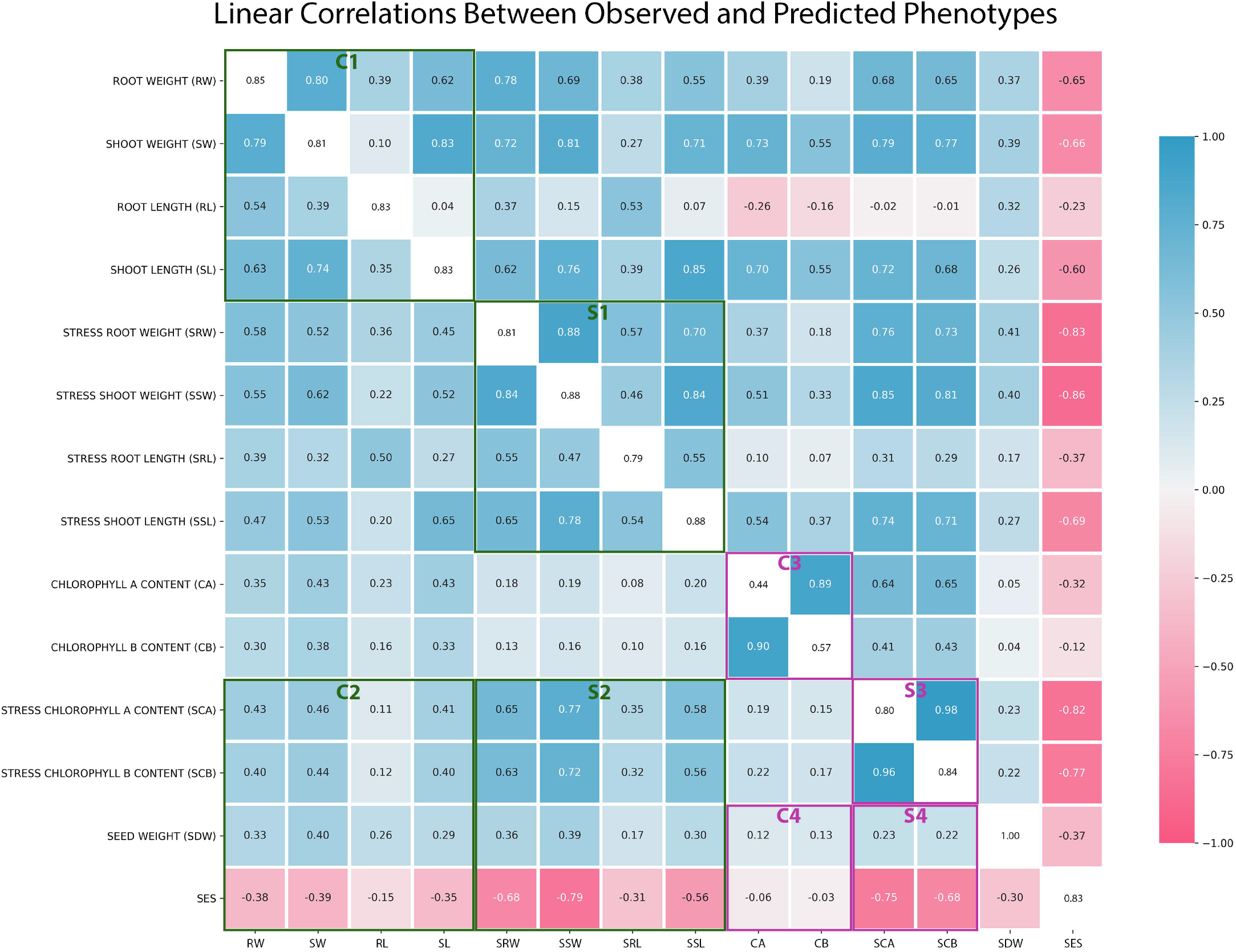
Linear correlations (r) between observed and predicted phenotypes. The bottom triangle of the white diagonal shows correlations between observed phenotype values and the top part is comprised of correlations between values estimated from genomic prediction. The white diagonal is the correlation between observed and predicted values for each phenotype. In the highlighted rectangles, C stands for control condition and S stands for stress condition. C1 to S1 and C2 to S2 depicts a fortification of linear correlation between biomass traits when salt stress is imposed. C3 to S3 show the same trend for chlorophyll traits. C2 to S2 and C4 to S4 further show that the enhancement of linear correlation brought about by salt stress applies to SES as well.

In figure 3, we observe fortification of trait correlations in salt stress conditions relative to control conditions. This is particularly true for biomass, chlorophyll and SES traits which implies that the encumbrance of salt stress on a plant prevents the disproportionate gain of biomass. Since plant biomass and chlorophyll content is markedly reduced under salt stress (supplementary table 3), we can conclude that salt stress does not affect all phenotypes for all genotypes at the same rate and the genetic advantages that any genotype has in terms of tissue growth (root/shoot) or chlorophyll accumulation will be compromised by the effect of abiotic stress at a greater magnitude than traits which have near-baseline values. On account of linear correlation, plant root length stands out to be the most independent phenotype, especially in control conditions.

Variables following a standard normal distribution are more statistically reliable and allow parametric statistics. Gaussian residuals are a pre-condition for linear modelling and regression-based association testing customary to advanced GWAS models. Density plots for all processed and derived phenotypes with subgroup specific curves for 82 Indica and 78 Aus varieties are tiled in supplementary figure 2. The biomass related traits we measured mostly followed a normal distribution with the exception of plant shoot length in both control and stress conditions. The ion content and chlorophyll content traits displayed large skews with few extreme values in either poles. Out of the ion content phenotypes, only shoot sodium content in stress condition exhibited a gaussian distribution (Supplementary figure 2). The seed property phenotypes were considerably normal, but influenced by extreme values. This could be anticipated as few traditional varieties exhibit particularly short and small seeds while some varieties have very long, bulky seeds. Filtering out few extreme values mended the majority of the distributions to fit a normal curve. Supplementary table 6 lists the Shapiro-Wilk statistics for all phenotypes prior to filtering and the number of data points that needed to be omitted to improve the distribution. Filtering data for any anticipatory means is bad statistical practice. The data filtration was done to examine the nature of the phenotypes as random variables and for parametric tests carried out post-GWAS. We did not use the filtered data in the primary association testing as it would compromise statistical power and risk spurious associations.

At this point, because of the presence of some highly non-normal traits (supplementary table 6), we looked towards data transformation methods. Traditional data transformation protocols for non-normal data often obscure the connotations of discovered significant associations [21]. The WarpedLMM package [21] learns an optimum transformation function from non-normal observations as it models the data to the corresponding genotypes. The module works as a wrapper on top of the previously published FaST-LMM module [34]. We have accordingly transformed all datapoints from all phenotypes assigned for association studies using WarpedLMM prior to model fitting in GWAS. The significant changes in the skew and range of the data distributions after transformation by WarpedLMM can be seen in figure 4. The untransformed phenotypes were scaled to allow visualization of all 39 phenotype densities in one plot. Scatter plots of residuals from genomic estimates of phenotypes calculated before and after transformation can be seen in supplementary figure 3. The transformation method improved the centered clustering of residuals around zero and their uniformity in dispersion.

**Figure 4.**
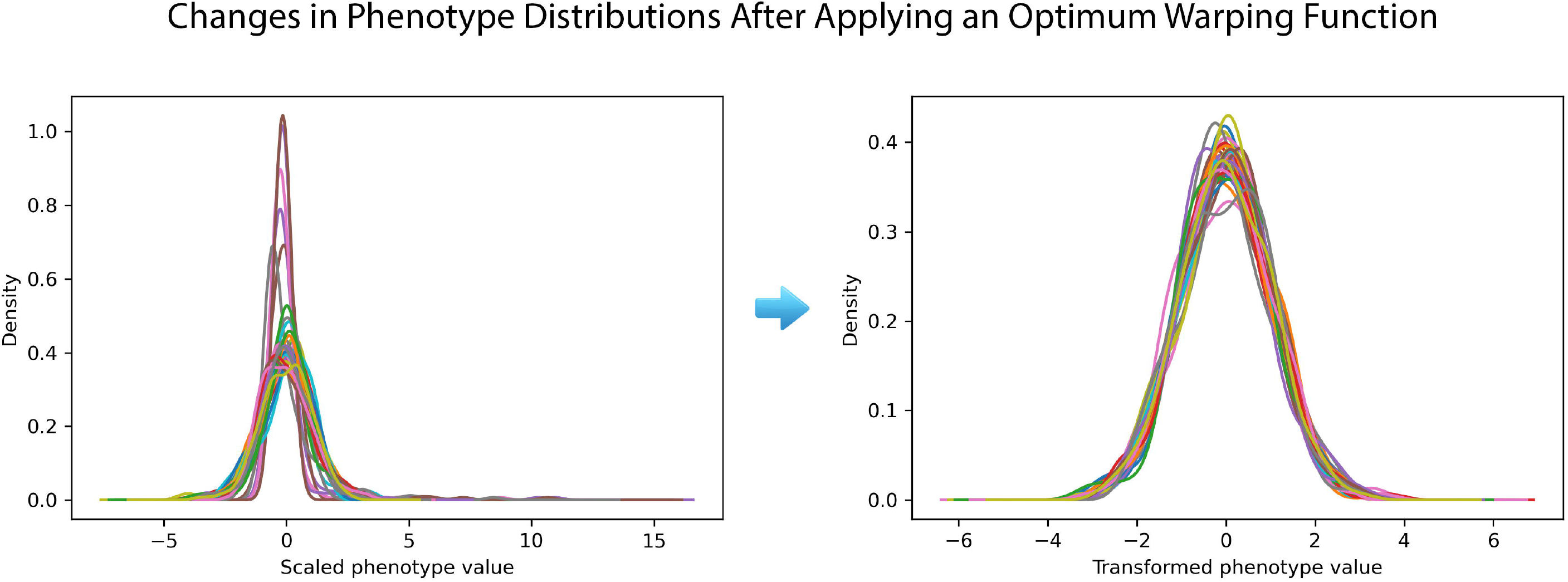
Changes in phenotype distributions after applying an optimum warping function. The plot on the left shows that densities of some the observed phenotypes were highly irregular even after scaling but when transformed using WarpedLMM they all appeared normal.

### 3.3 Extensive genome-wide association studies

GWAS results discussed in this study were performed by BLINK [26]. The BLINK algorithm [26] implemented in the GAPIT software package [27] is a recent innovation for powerful statistical association testing at considerably lower computational cost. BLINK achieves enhanced computational time by employing a linkage disequilibrium (LD) based iterative marker pruning strategy that fits two fixed effect models with a filtering step involving the second model. The algorithm is discussed in detail in the methods section. We also fit a more commonly used compressed mixed linear model (CMLM) to our phenotypes with optimum compression level to compare and confirm significant associations. Three independent studies on the whole population and Aus and Indica subpopulations were carried out respectively. 39 individual Manhattan and QQ plots each from three independent sets association tests are stored in the git repository under the ‘GWAS’ folder name. Table 2 lists 17 total QTLs identified and grouped into biomass, seed property or salt tolerance categories on the basis of their significant association to multiple traits belonging to the mentioned trait class. The significance threshold for multiple testing was set at -log(p) = 6.5 for the whole population and 6.4 for the two subpopulations. A common suggestive threshold of -log(p) = 5 was also set for all cases. The first identified QTL, qCDP1, was observed in plant shoot length measured at stress conditions in the Indica subpopulation (Figure 5 (a)). A suggestive peak on the same loci is found in the Manhattan plot for SES in the same subpopulation (Supplementary figure 4). The suggestive association found in SES made us consider role for qCDP1 in stress response. Two other associations noted for biomass traits were qCDP15 and qCDP17. Like qCDP1, qCDP17 can also be found associated with stress tolerance traits (Supplementary figure 4). qCDP2 is first observed in seed weight (Figure 6 (a)), but also found in seed height and seed volume traits (Supplementary figure 5). Three more seed trait loci, qCDP4, qCDP5 and qCDP14 are found significant in seed traits for Indica and the whole population. All four loci are suggestive in other seed related traits (Supplementary figure 4). qCDP3 relates to potassium content in plant shoots (Figure 7 (a)). Because of the central role of potassium and other cations in salinity tolerance, we observe qCDP3 as a stress tolerance trait. Four more signals, qCDP6, qCDP7, qCDP9 and qCDP10, relating to the effect of salinity on plant shoots shown in figure 7 (b) and (c) were discovered. qCDP8, qCDP11 and qCDP12 concerning root and biomass traits are observed in Figure 8. Figure 9 shows that apart from qCDP3, two more loci, qCDP13 and qCDP16 are associated with ion content phenotypes in stress conditions. 12 out of 17 QTLs in our findings were previously discovered by GWAS studies with similar phenotype connotations (Table 2). An extended version of table 2 with peak marker locations, original phenotype names and references [10, 11, 35-48] can be found in supplementary table 7.

**Table 2:**
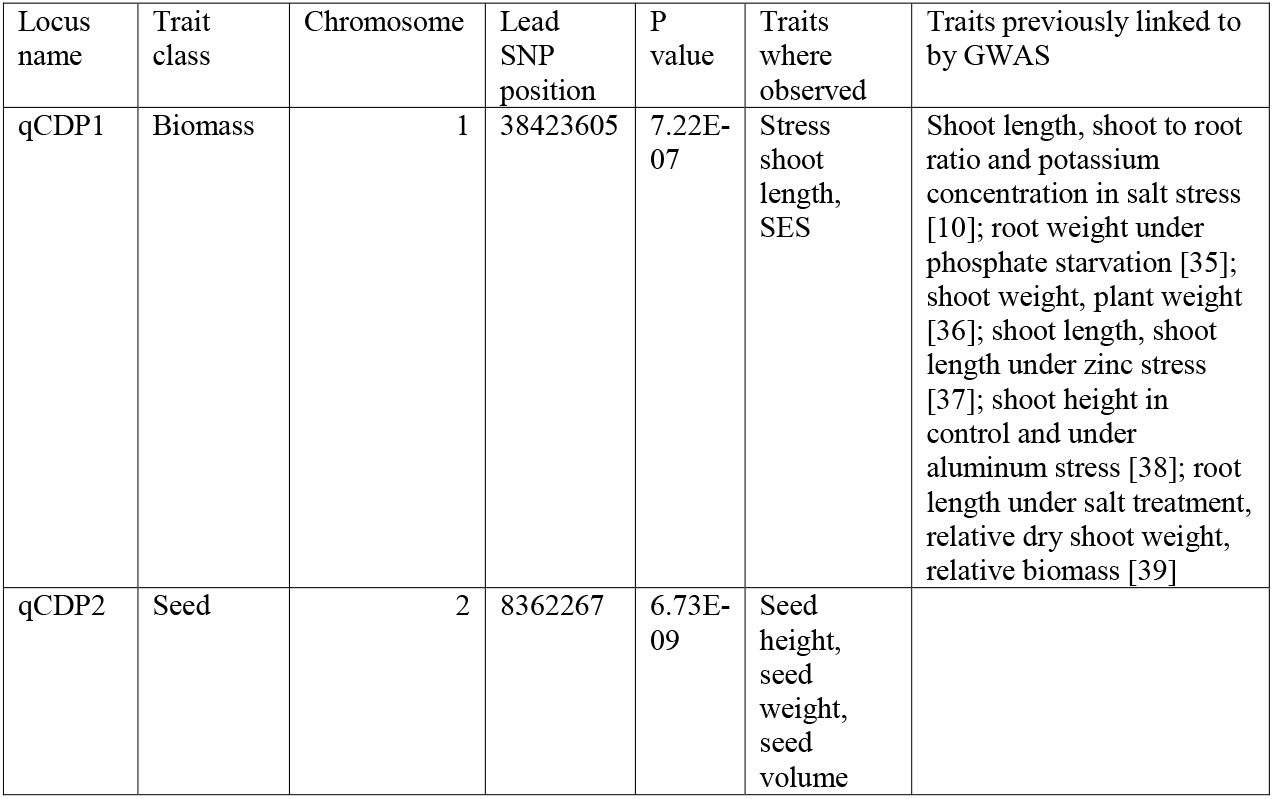
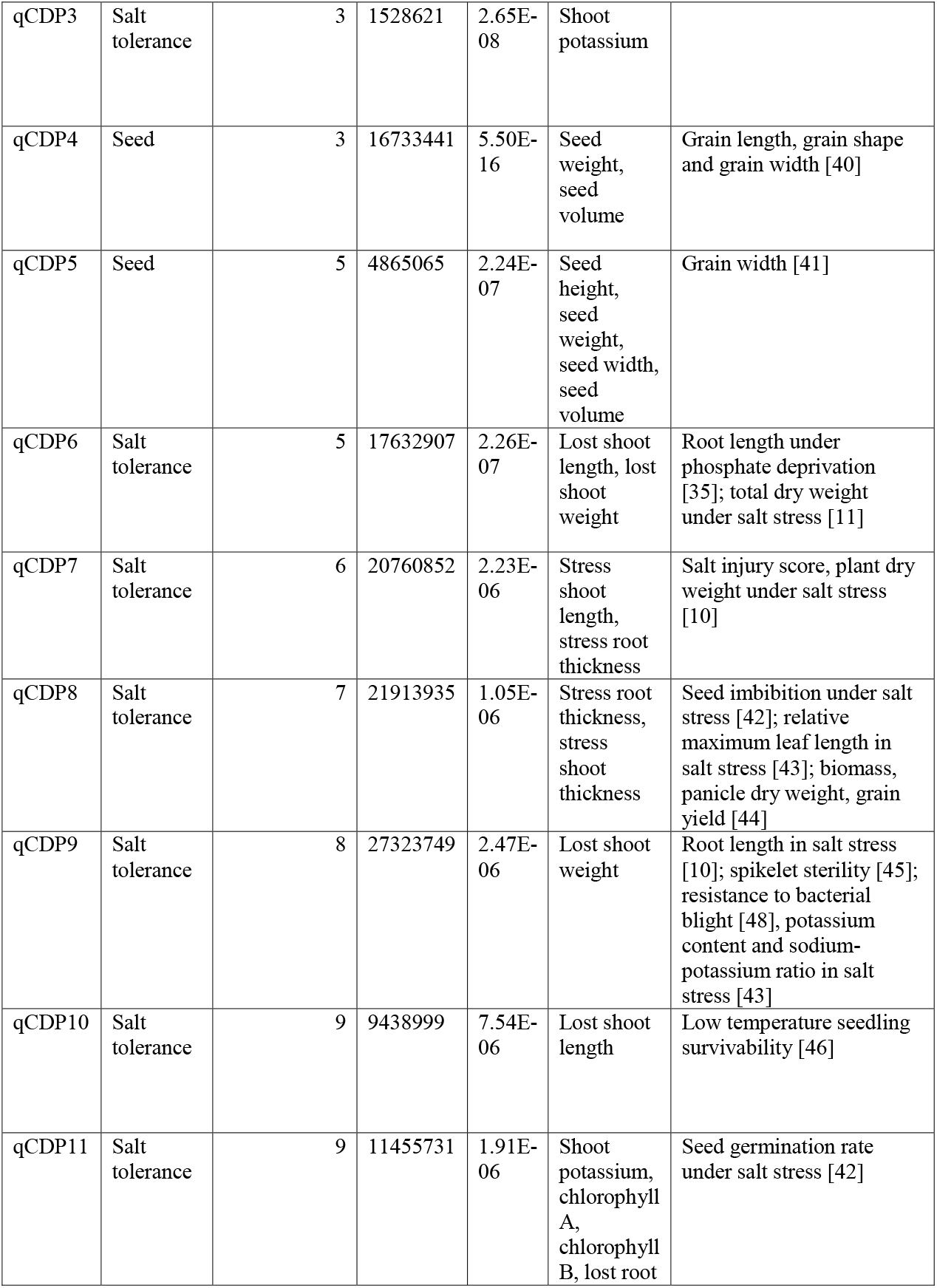
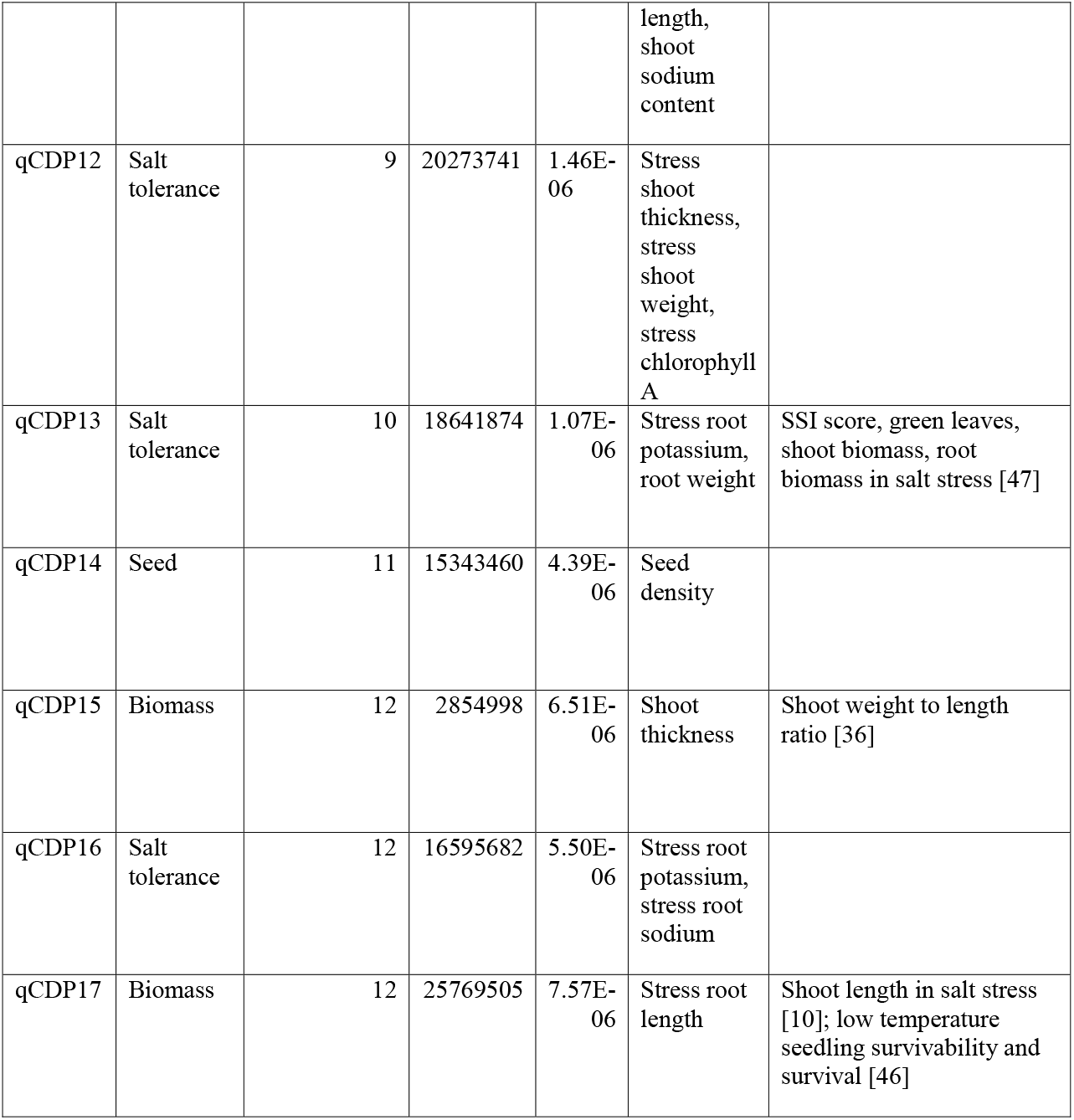
Identified Major QTLs.

**Figure 5.**
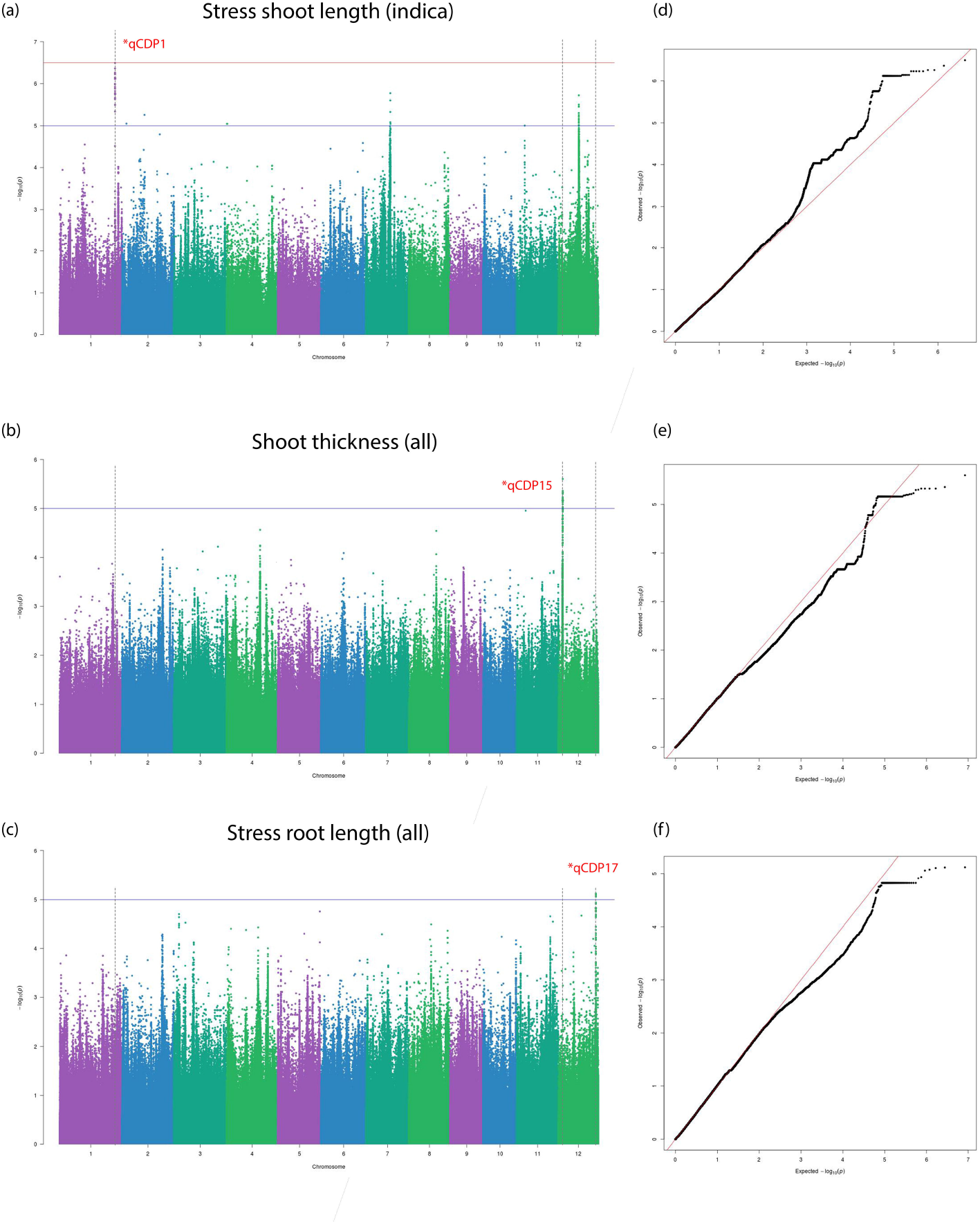
Manhattan and Q-Q plots for qCDP1, qCDP15 and qCDP17. Horizontal red and blue lines show the significant and suggestive thresholds respectively. Vertical black dotted lines show the locations of the three QTLs in the three Manhattan plots.

**Figure 6.**
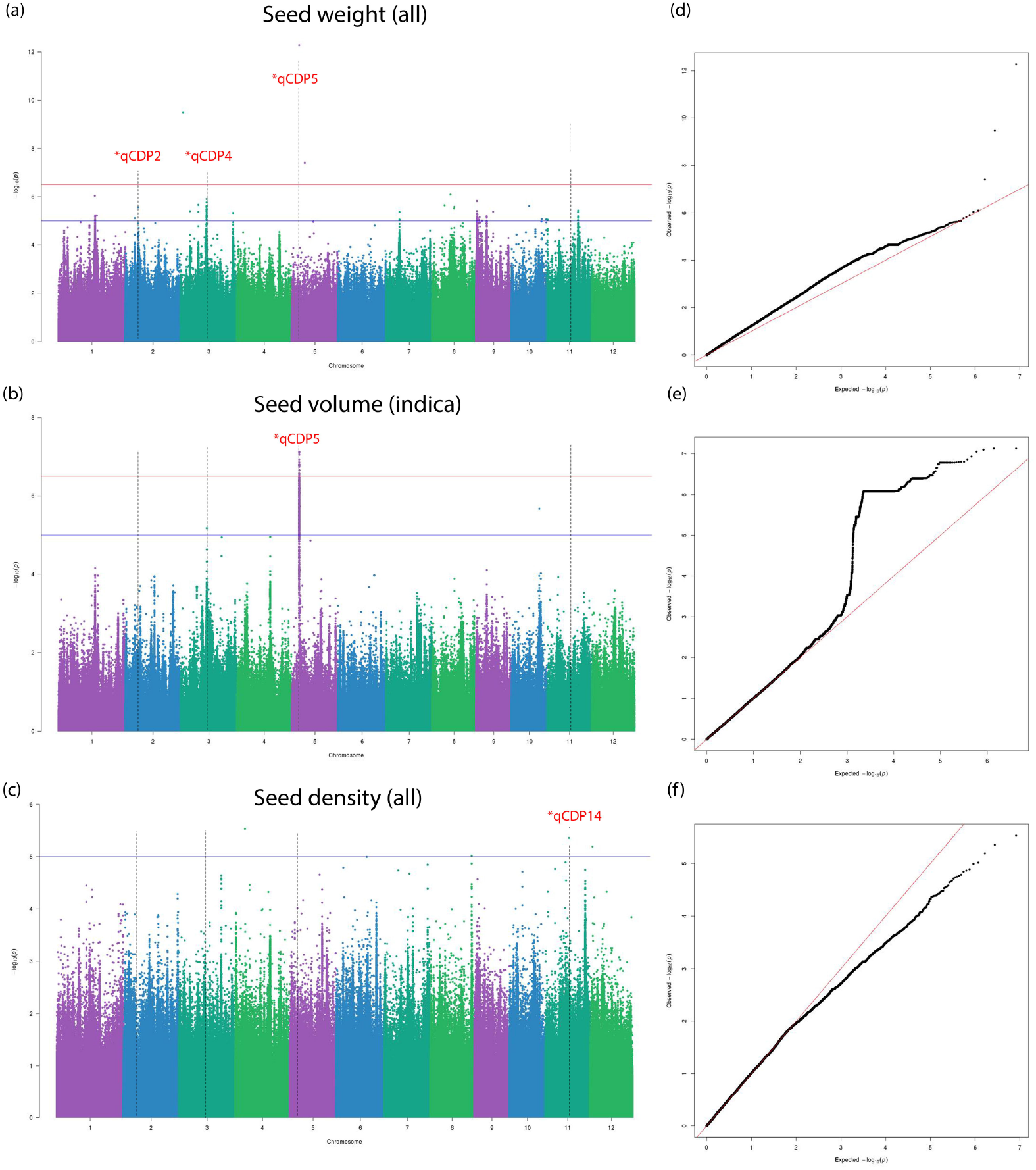
Manhattan and Q-Q plots for qCDP2, qCDP4 and qCDP5. Horizontal red and blue lines show the significant and suggestive thresholds respectively. Vertical black dotted lines show the locations of the three QTLs in the three Manhattan plots.

**Figure 7.**
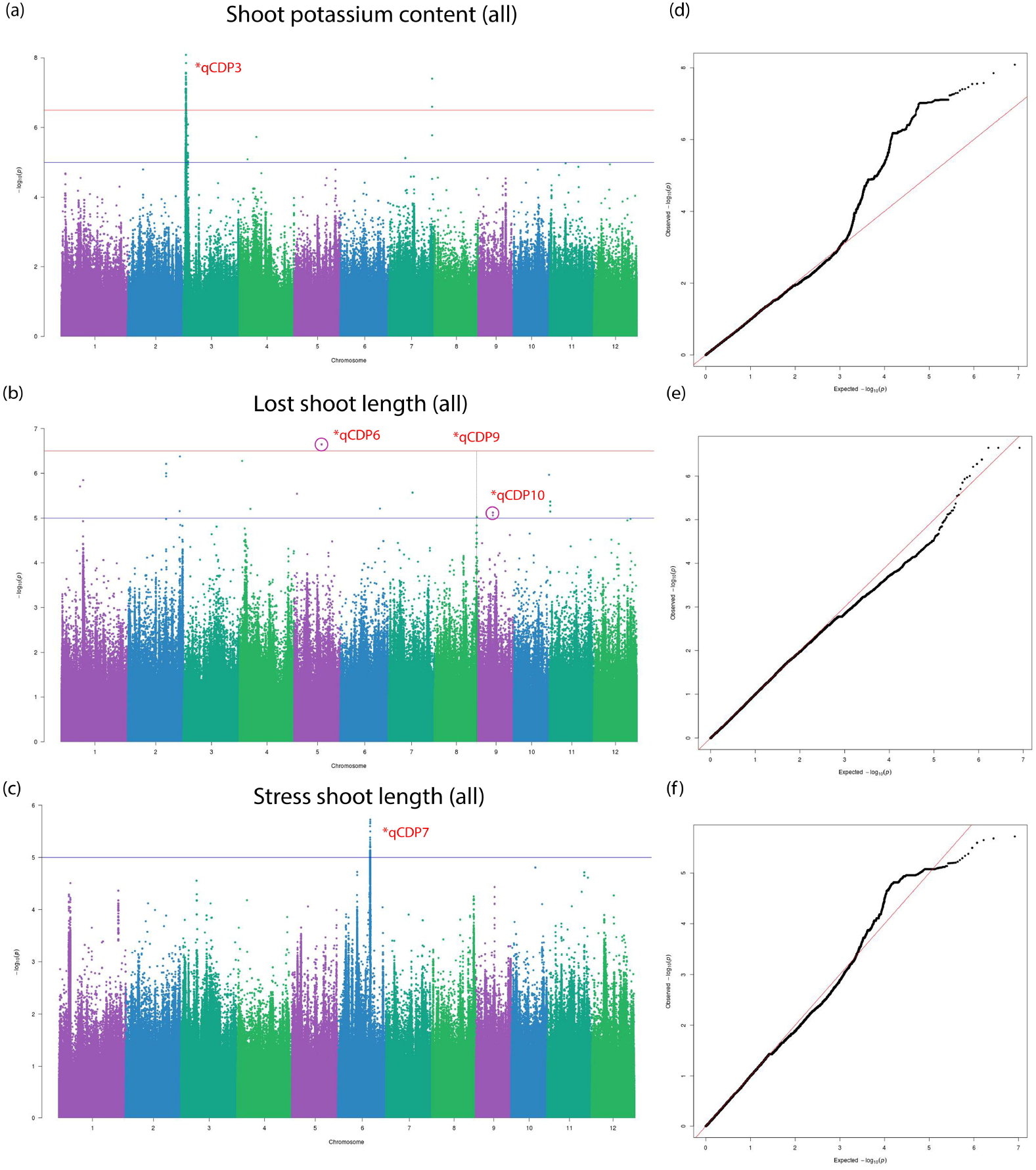
Manhattan and Q-Q plots for qCDP3, qCDP6, qCDP7, qCDP9 and qCDP10. Horizontal red and blue lines show the significant and suggestive thresholds respectively. Vertical black dotted lines show the locations of the five QTLs in the three Manhattan plots.

**Figure 8.**
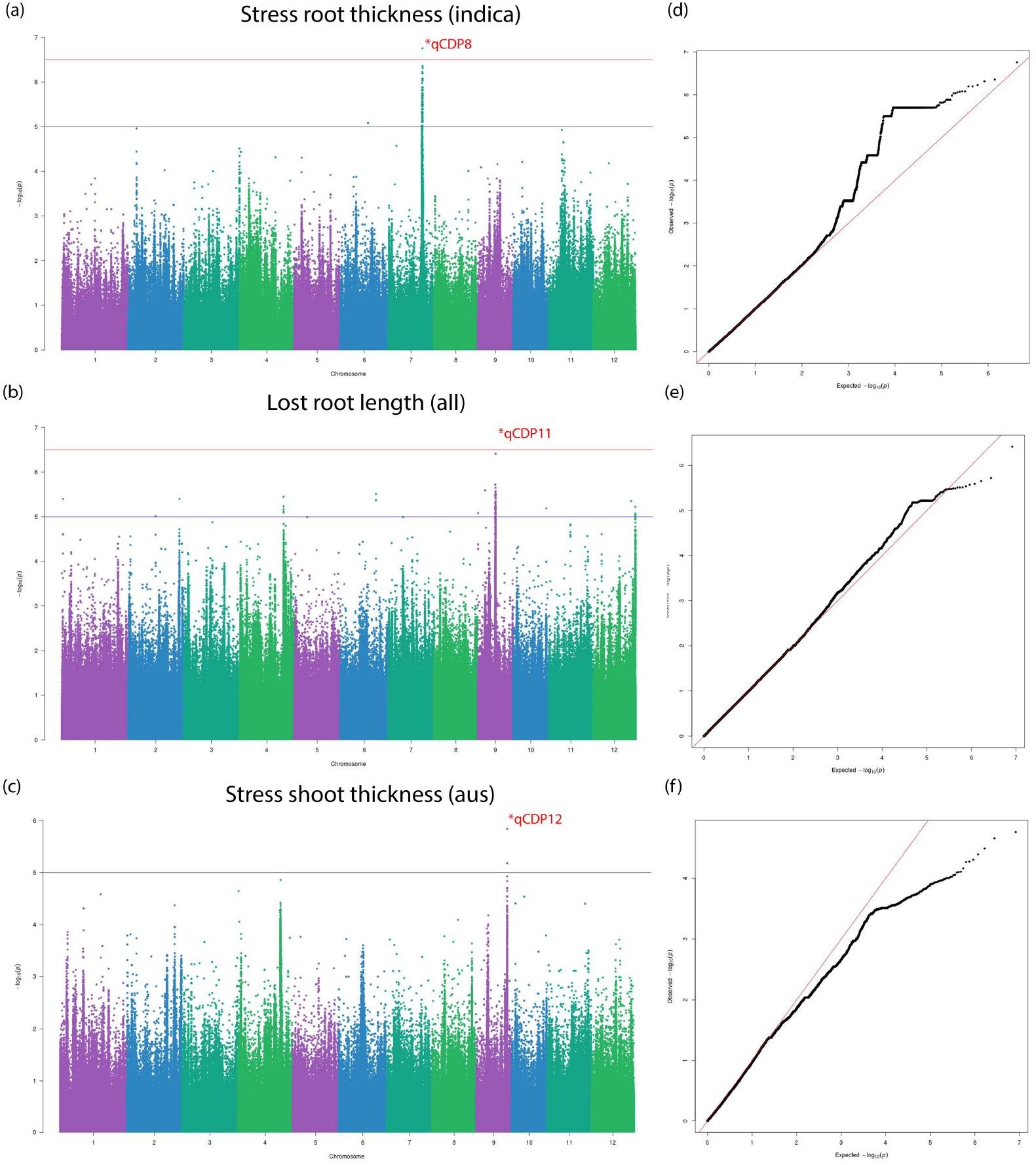
Manhattan and Q-Q plots for qCDP8, qCDP10 and qCDP12. Horizontal red and blue lines show the significant and suggestive thresholds respectively. Vertical black dotted lines show the locations of the three QTLs in the three Manhattan plots.

**Figure 9.**
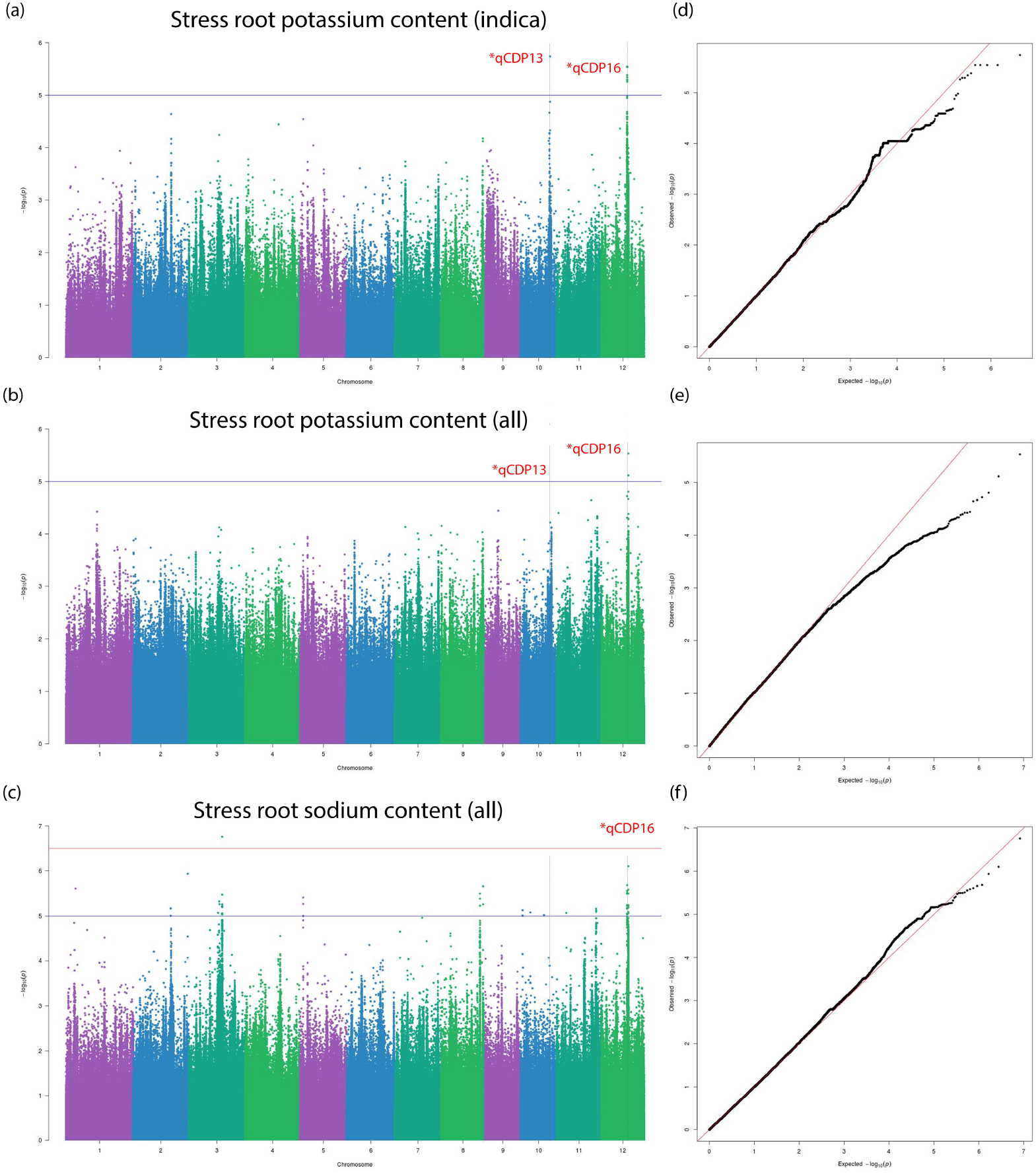
Manhattan and Q-Q plots for qCDP13 and qCDP16. Horizontal red and blue lines show the significant and suggestive thresholds respectively. Vertical black dotted lines show the locations of the two QTLs in the three Manhattan plots.

### 3.4 QTL mapping

The extent of linkage disequilibrium in rice has been found consistently in the range of 100kb to 200kb [41], although some authors suggest that it could extend over 500kb for some subpopulations [49]. In our experiments, LD decay to half maximal value of r^2^ was observed at a distance of ∼70kb in the whole population, ∼100kb in the Aus subpopulation and ∼50kb in the Indica subpopulation (Supplementary figure 6). Based on these assumptions, we defined the range of our QTLs to be from 100kb upstream of the peak SNP up to 100kb downstream for association mapping. 399 markers showing significant and suggestive associations within the sequence range of known rice genes are described in supplementary table 8 with metadata from the RAP-DB database [13]. All significant and suggestive markers from validation tests using the CMLM model can be found in supplementary table 9. We reviewed the annotations for genes situated in the vicinity of these significant associations for the elucidation of functional roles in regards to the implied phenotypes. From these annotations, 46 genes with notable functional roles corresponding to their associated phenotypes are included in table 3.

**Table 3:**
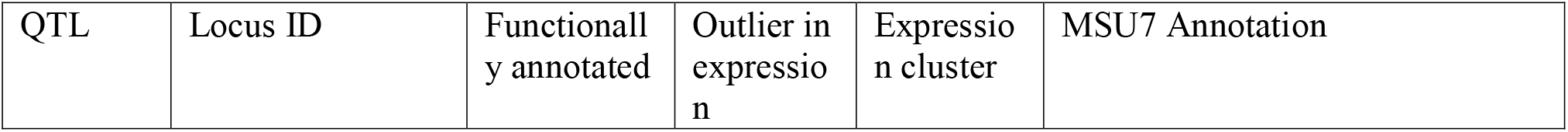
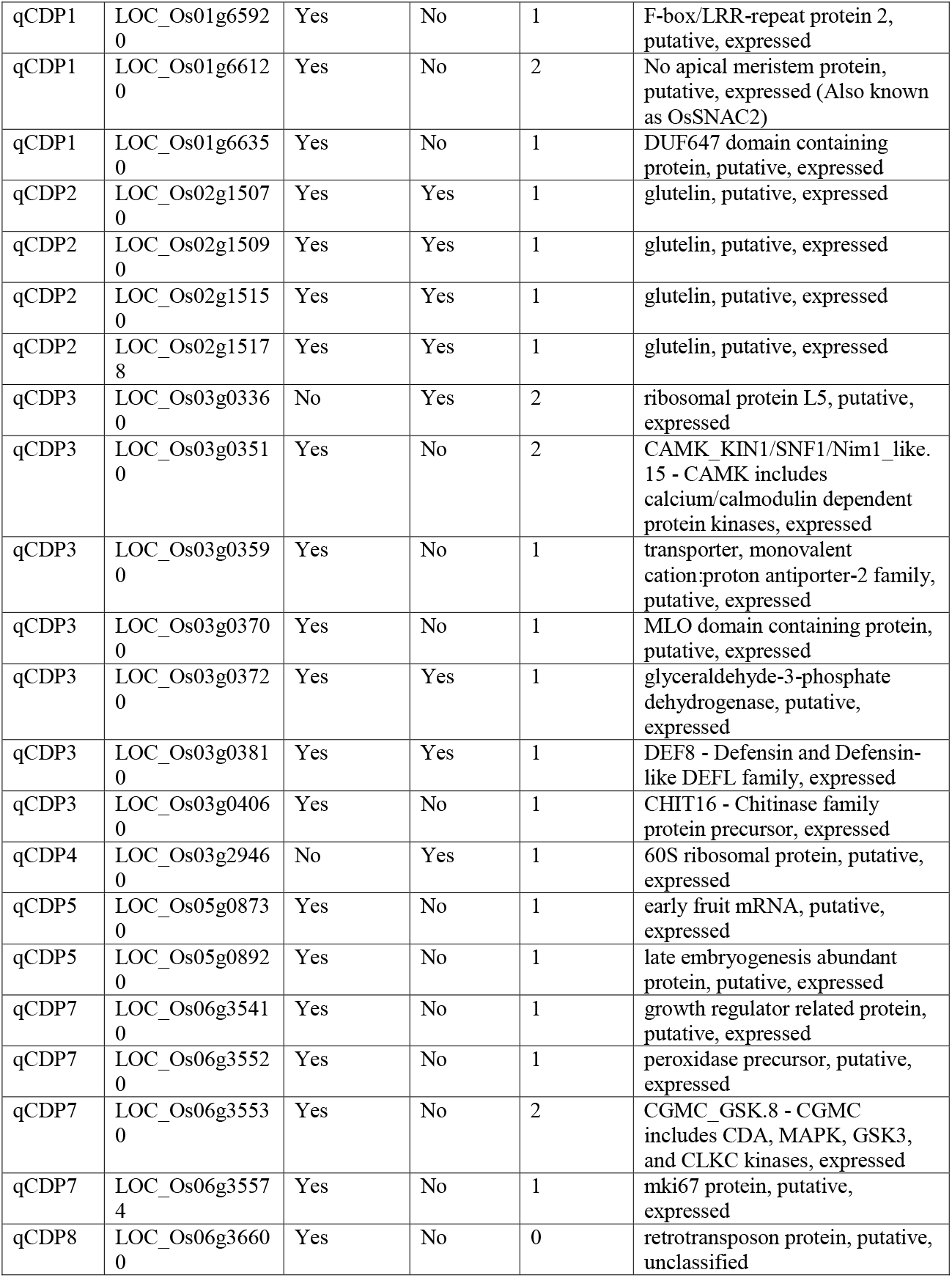
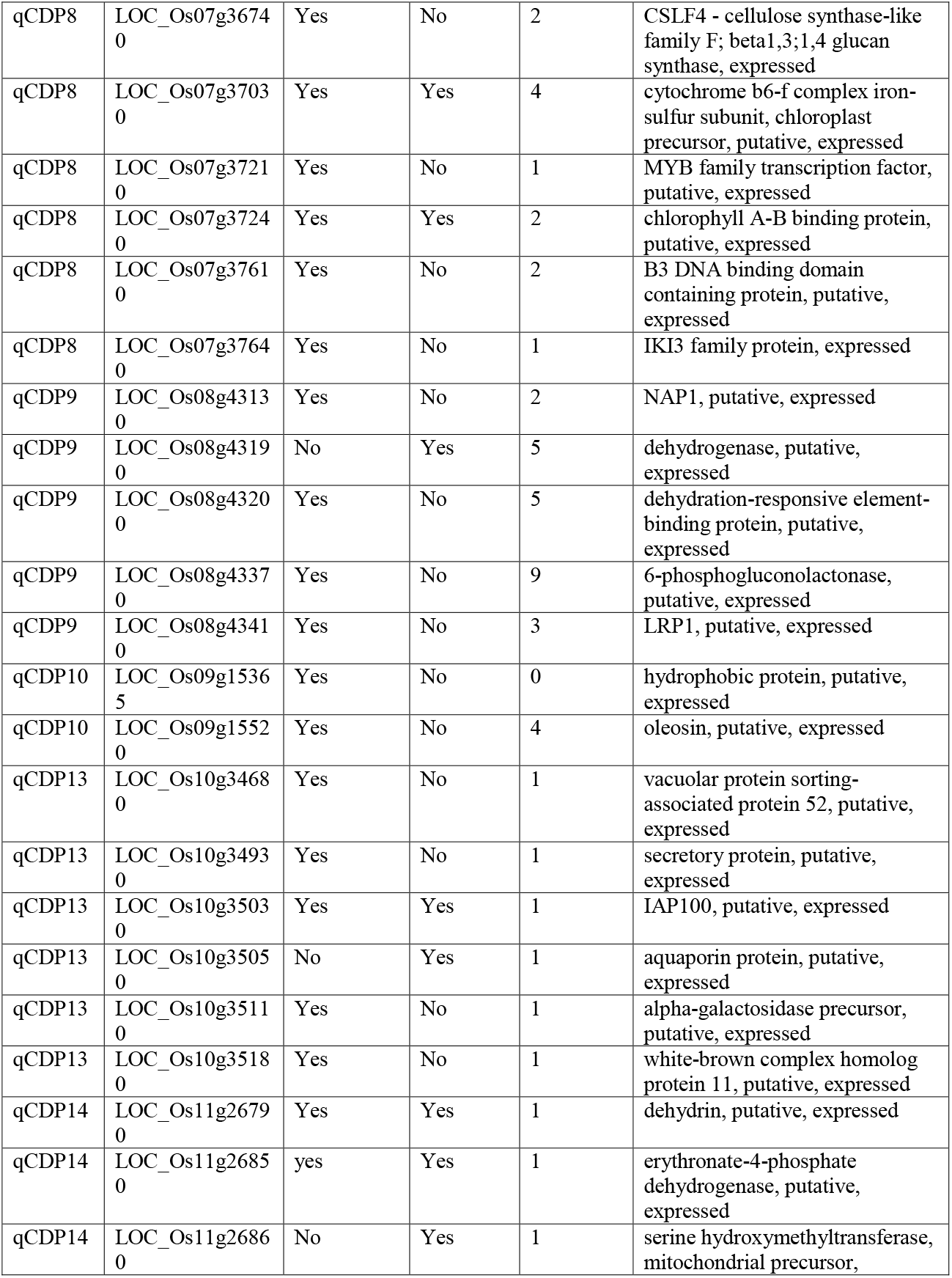
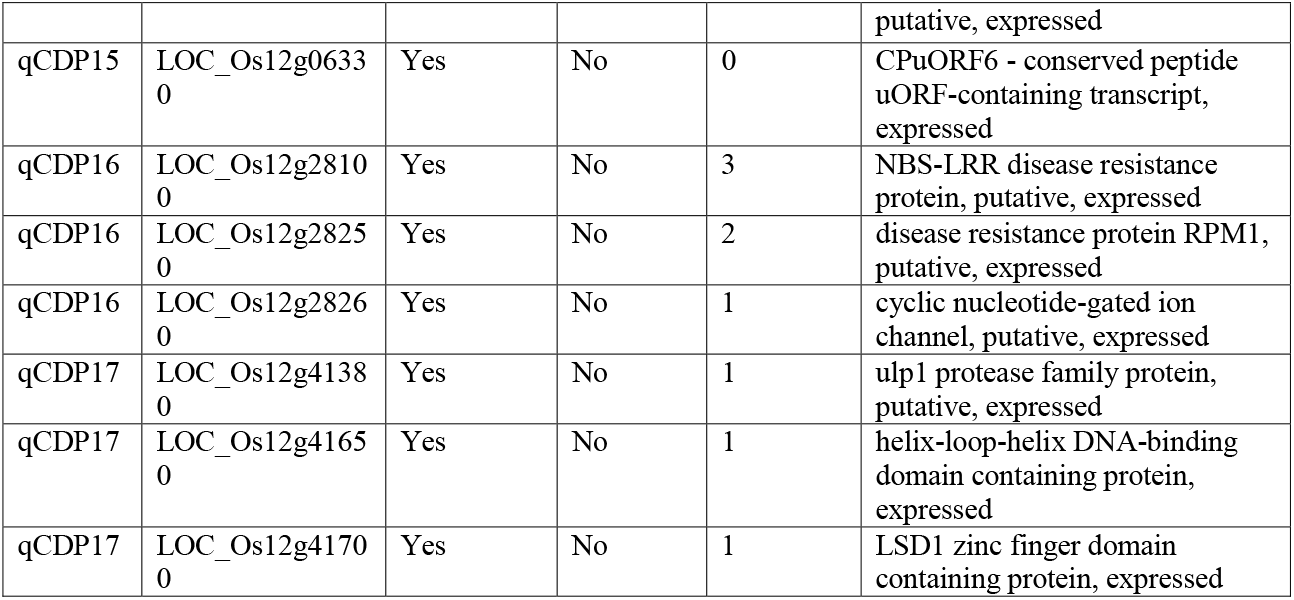
Reviewed candidate genes.

The most significant marker association, with a -log(p) value of 15.26 was observed for seed length and directly maps to LK3 (RAP-DB ID: Os03g0407400), a gene named for its functional role in the regulation of grain size. Another highly significant marker for seed height was found to be OsEFCAX2 (MSU7 ID: LOC_Os02g14980), which is a calcium transporter expressed in all tissues having 4-fold greater expression in pollinated embryo. One significant marker from qCDP3 associated with shoot potassium content could be mapped to OsGSL10 (MSU ID: LOC_Os03g03610), a metabolic enzyme having 2-fold higher expression in shoots than any other tissue. In the same QTL, another cation antiporter associated to shoot potassium content, LOC_Os03g03590, could be mapped by a highly significant marker. This gene is also ubiquitously expressed with significantly higher expression in leaves and shoots. Based on its peptide sequence, the transporter protein is annotated to be localized in the chloroplast (Uniprot ID: Q10SB9). Within the same locus, two notable genes: OsCP29 (MSU ID: LOC_Os07g37240) and OsPetC (MSU ID: LOC_Os07g37030), are also localized in the chloroplast. Both these genes are expressed all throughout the plant in markedly higher expression levels and have almost 10-fold higher expression in leaf tissues. The presence of a chloroplast localized cation transporter adjacent to two particularly important chloroplast related genes in a gene locus identified through shoot potassium content imparts sufficient prospect to the role of this transporter protein and chloroplasts in the regulation of potassium ion content in plants. From the MSU7 brief loci annotations, genomic copies of the OsNAC6 gene (MSU ID: LOC_Os01g66120) which is known to have a functional role in plant development and stress tolerance, could be mapped to 5 identified QTLs: qCDP1 (LOC_Os01g66120), qCDP2 (LOC_Os02g15340), qCDP3 (LOC_Os03g03540), qCDP15 (LOC_Os12g05990) and qCDP17 (LOC_Os12g41680). qCDP1, qCDP15 and qCDP17 are QTLs belonging to our biomass category (Table 2) and qCDP2 is related to seed property. This is good evidence to suggest that OsNAC6 has a prominent role in embryogenesis, germination and biomass gain in rice. The gene has also been implicated in stress response and abiotic stress tolerance which holds to its presence in the salt stress related QTLs: qCDP1, qCDP3 and qCDP17.

### 3.5 Gene expression profiling and candidate gene selection

We selected 46 candidate genes in table 3 for further analysis based on review of available gene annotations. There are drawbacks to selecting candidate genes based entirely on metadata. These include the lack of curation and questionable quality of some few annotations. A greater concern is the inclusion of pseudogenes, gene fragments or genes lacking the appropriate cis-acting elements needed to enable the biological information pathway. Thus, to select a set of informative candidate genes with authentic functional roles, we collected plant tissue-specific gene expression data from the MSU7 database [13]. We visually profiled the expression signature of all genes located within the defined range of the 17 identified QTLs. To remove very sparsely expressed genes and reduce skewness, the expression table was filtered under the criteria that every row sum value is greater than 1 and log transformed by the function f(x) = log_2_(x+ 1) for ease of visualization. A total of 340 candidate genes from within the 17 QTLs that remained after pruning of poorly expressed genes by their raw FPKM read counts can be found in supplementary table 10. We sorted the genes first by QTLs followed by trait class and plotted the expression table as a heatmap seen in figure 10 (a). Five genes were found to have very high relative expression levels. These five genes are included in table 3 as outliers in gene expression making the total candidate gene count 51 in the table. In the expression heatmap, genes belonging to the seed trait QTLs are found to have noticeably higher expression in seeds after pollination. The genes associated with stress tolerance are markedly more expressed across all tissues and have particularly higher expression in the seedlings. Because the stress tolerance related candidate genes show overall uniform expression levels in figure 10 (a), a cluster map for these genes has been plotted in figure 10 (b). The salinity tolerance related genes could be designated to 4 expression groups and 10 sub-clusters based on their expression levels in different tissues (Figure 10 (b)). To group candidate genes by their inter-tissue expression signature, we clustered genes having similar gene expression profiles into 10 groups by hierarchical clustering. Genes belonging to the same cluster will display similar relative expression patterns across all tissues. Details on every selected candidate gene and the expression cluster they belong to can be found in supplementary table 11.

**Figure 10.**
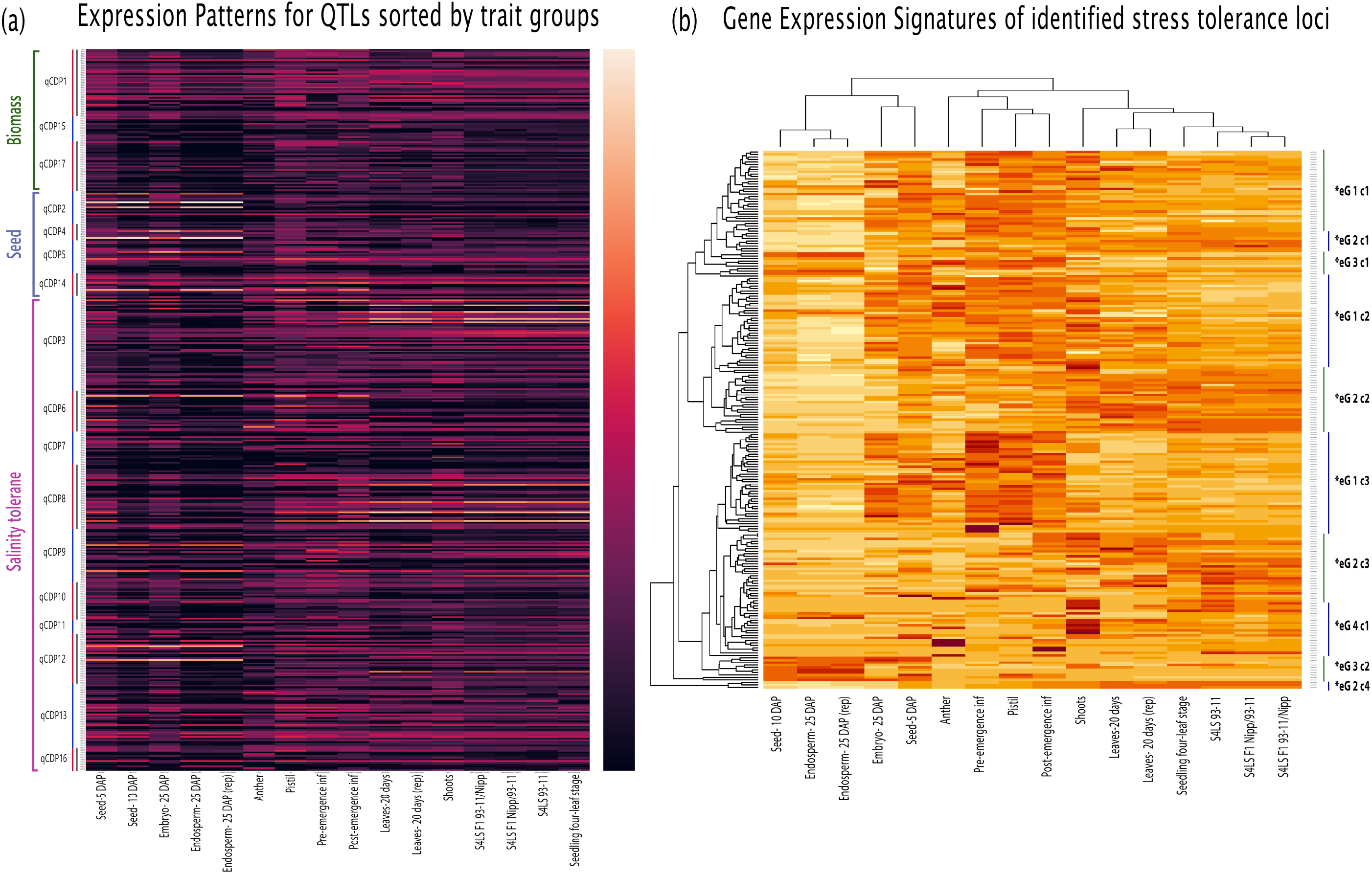
Gene expression heatmaps for 340 genes from the 17 identified QTLs. (a) Expression patterns for QTLs sorted by trait groups. Vertically, genes are first arranged by QTL and then by trait class. Biomass related genes are found to be expressed more in mature tissues than seed and pollinated embryos. Some seed related genes have visibly high expression in seeds and embryos. Genes belonging to QTLs tagged by salinity tolerance display a more uniform expression pattern with relatively high expression in mature tissues. (b) A clustered heatmap of 211 genes from supplementary table 10 implicated in salinity tolerance. Genes are clustered into four expression groups denoted *eG1-4 from visual inspection and these groups appear as 10 separate clusters shown using c1-4.

### 3.6 Haplotype testing and candidate gene curation

Out of all SNP markers, we extracted SNPs including indels that occur within the 340 candidate genes. According to MSU7 annotations, the whole set of candidate genes are transcribed to 427 unique and expressed mRNA transcripts. 28188 possible marker-transcript pairs within candidate gene regions were determined, and the consequence of the substitution at the polymorphic allele was elucidated. The complete table of SNP effects including the aligned transcript, site of mutation (exon/intron), occurrence of frameshift and consequence of substitution (synonymous, missense, nonsense) is provided in supplementary table 12.

For each substitution, we tested for significant difference of means between the observed genotypes using One-way ANOVA and the student’s t-test. In One-way ANOVA, three genotype groups were denoted AA, Aa and aa defined as those having two, one and no substituted alleles respectively and for the student’s t-test, only homozygotes were tested. Here, we ignore the confounding caused by population structure and carry out general hypothesis testing as these candidate genes all belong to QTLs that were originally identified using protocols that adjust for confounding with necessary corrections. Since the presence of extreme observations could bias the results for one genotype group, instead of using all the observed phenotype values, we used a filtered subset of values for each phenotype from the information previously presented in supplementary table 6. FDR correction for multiple testing was conducted by the Benjamini-Hochberg method. The substitutions showing a significant difference in the means of their observed phenotypes at an FDR adjusted p value cutoff of 0.01 along with their corresponding phenotype effects can be found in supplementary tables 13 and 14 for One-way ANOVA and the student’s t-test respectively. QQ-plots for p values from One-way ANOVA and the t-test are shown in supplementary figure 7.

It is understood that the consequence of most intronic and nonsynonymous mutations is trivial, having little to no impact on protein function. Some intronic mutations can have effects from altering splicing sites, but the frequency of such an event is rare compared to the large number intronic polymorphisms that are observed. Other nonsynonymous mutations may change the rate of transcription and translation by affecting the interactions of transcription factors and DNA binding proteins. Transcripts having nonsense codons in the exon region are more likely to be functionally impaired but only variants having frameshift mutations that are not towards the very end of the coding sequence are certain to have lost some biological function. Under this model of thought, we sought to identify single polymorphisms having the most consequential functional effect at the protein level. Out of 28188 substitution events within 17 QTLs, we found 33 total frameshift and nonsense mutations. The summary statistics for these 33 SNPs belonging to 21 genes along with traits where significant differences were observed is presented in table 4. We impart novel functional roles to these 21 impaired genes compiled in table 5 on the basis of gene locus, expression signature, LD-based haplotype conservation, previous studies and significant differences in phenotype when prematurely truncated. For each assigned function, individual scores are given at every evidence level. The given QTL evidence score is 0 when the assigned function is not supported by any trait associations, 1 when supported by one significant trait association and 2 when supported by more than one trait association (table 2). Expression scores can be 0, 1 or 2 based on whether a transcript has relatively low expression, relatively high expression or very high expression in relevant tissues (supplementary table 10) respectively. A literature score of 1 is given when the suggested function is implicated in gene annotations or ontology terms and for markers whose LD value is greater than the mean LD value of all markers within the QTL (table 4), a haplotype conservation score of 1 is given. A confidence score for the given annotation is obtained from the sum of these scores.

**Table 4:**
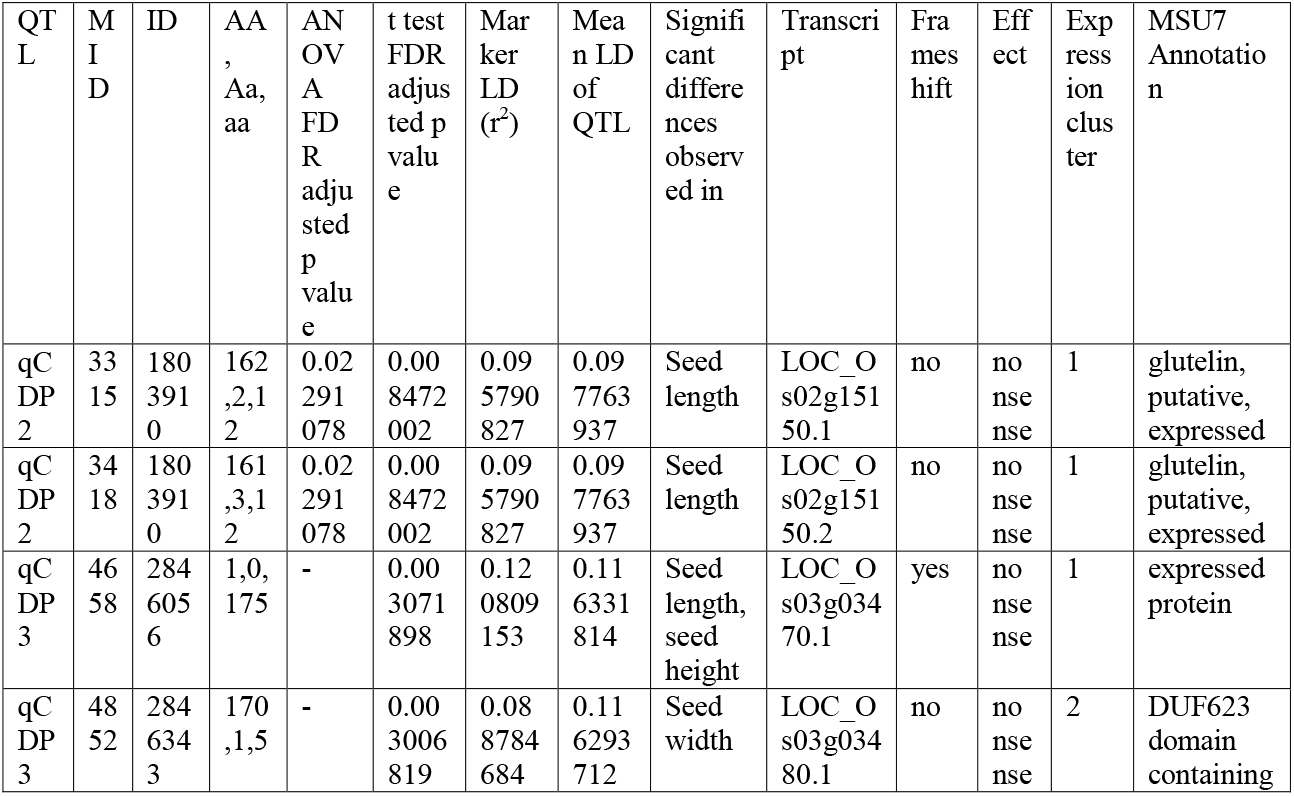
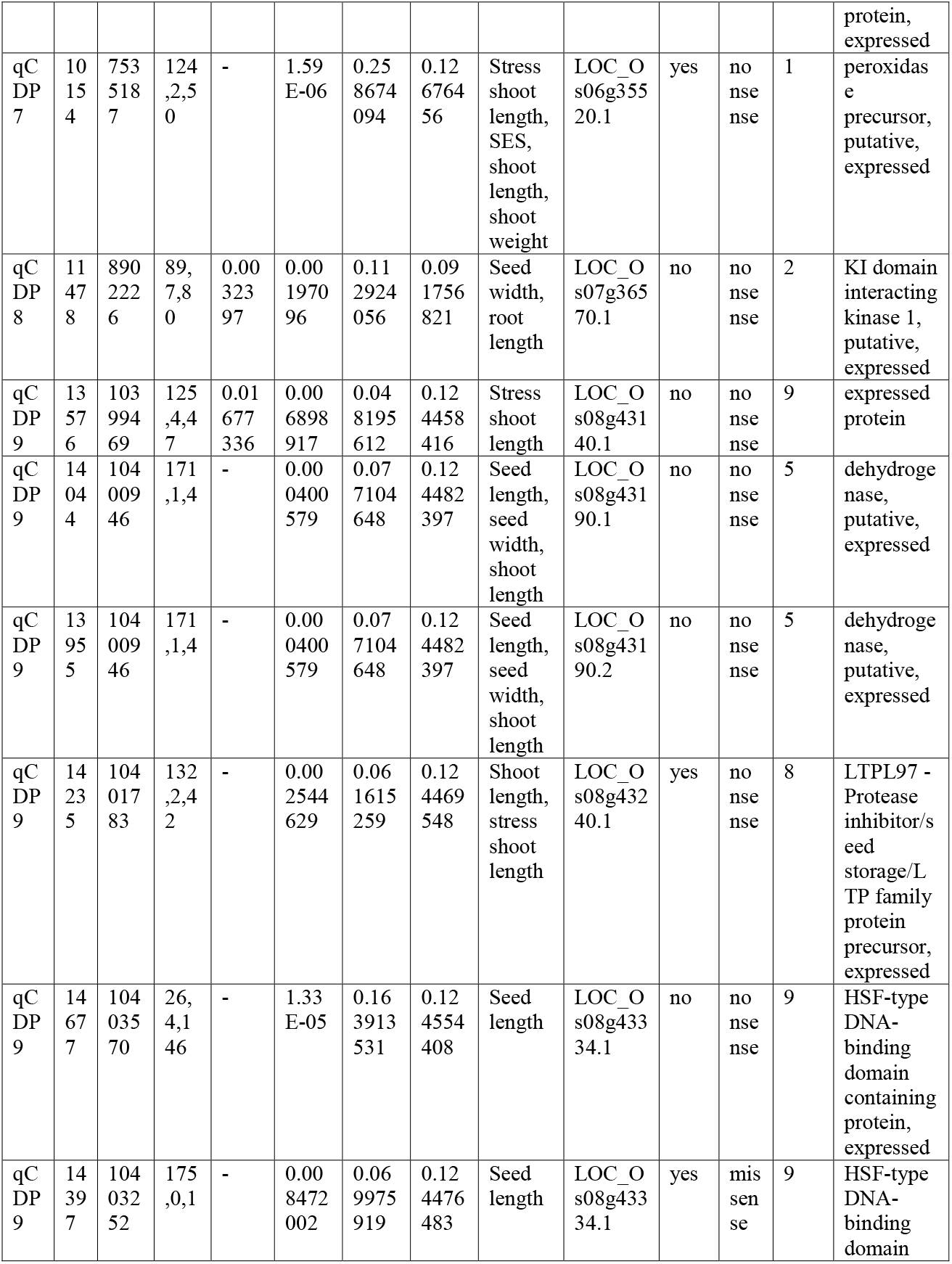
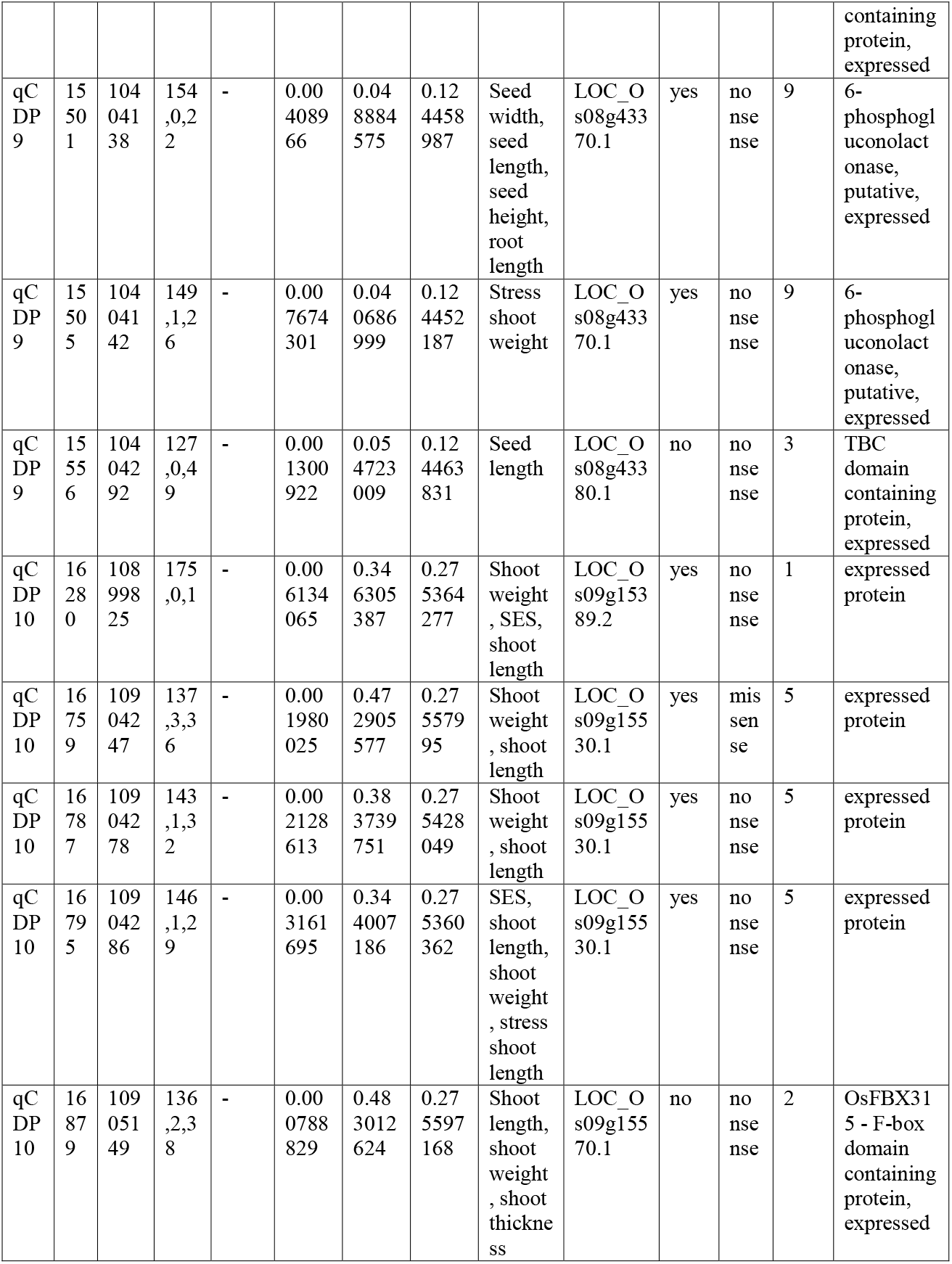
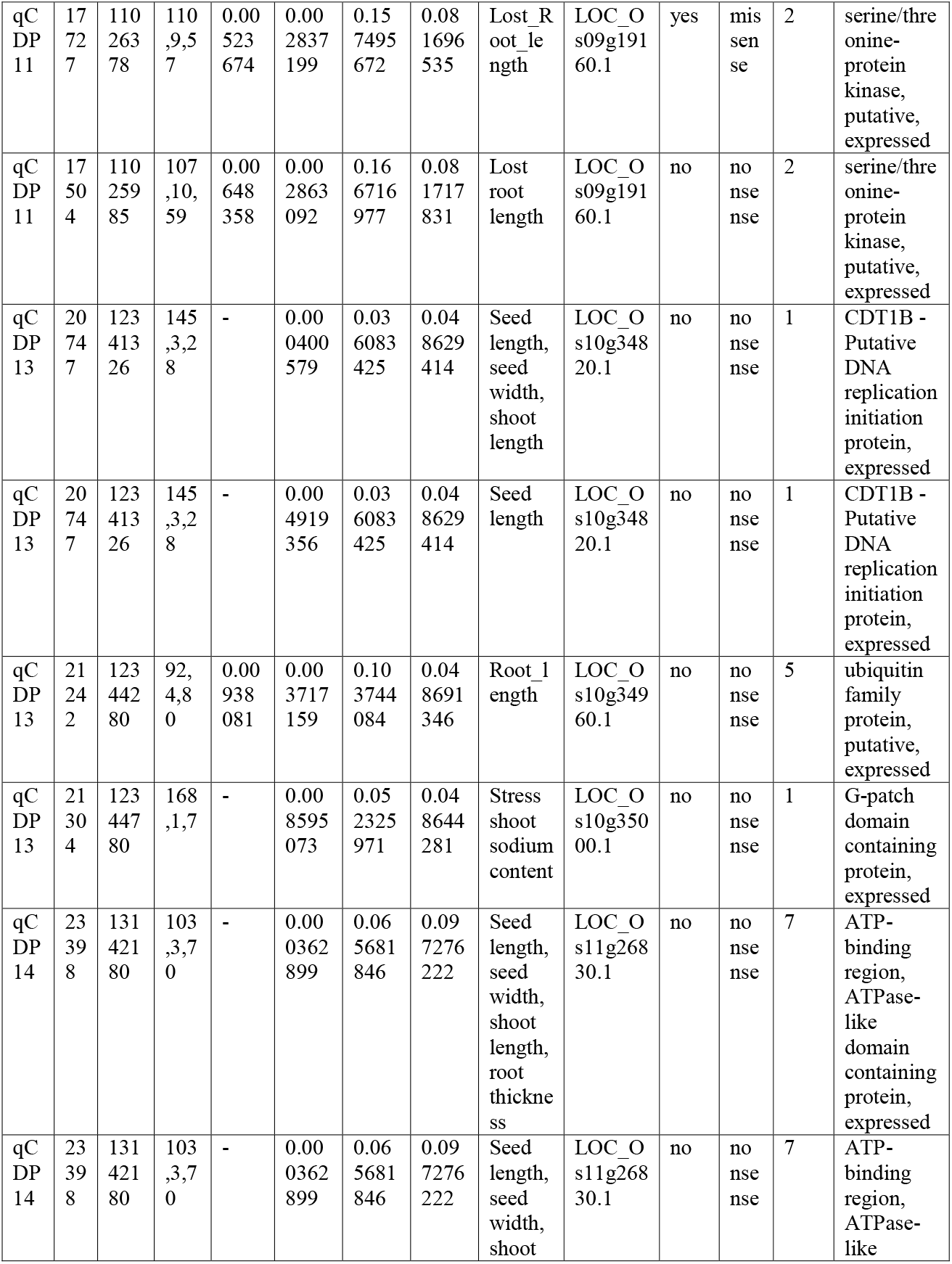
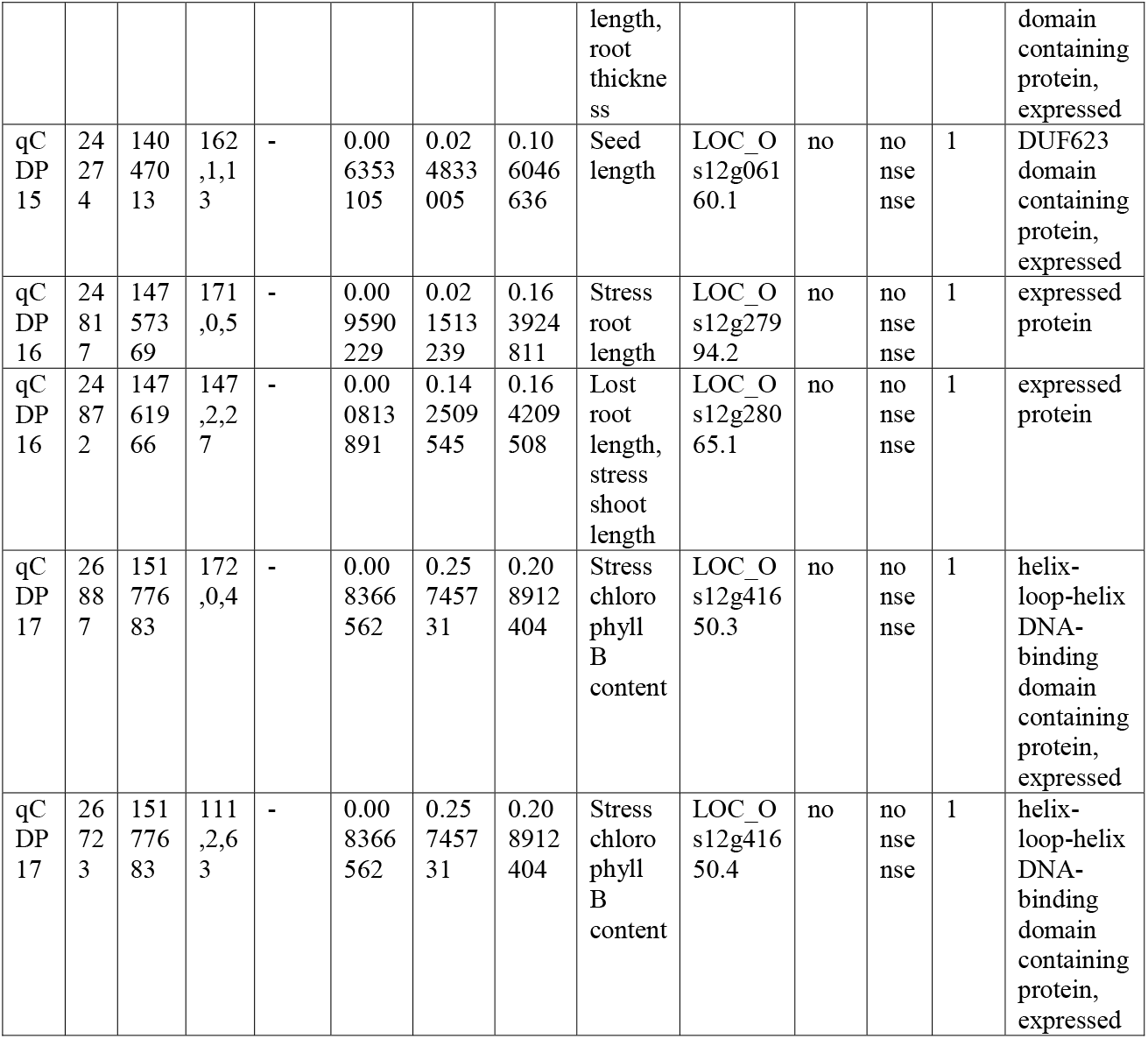
Frameshift and nonsense mutants.

**Table 5:**
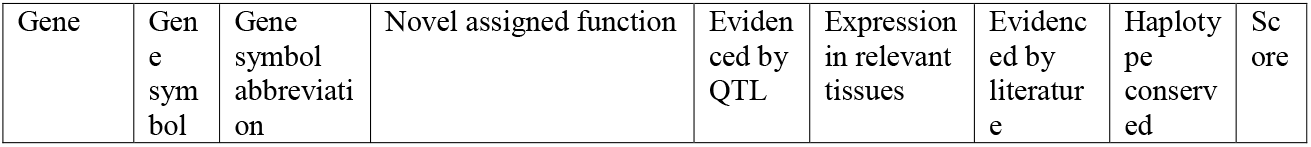
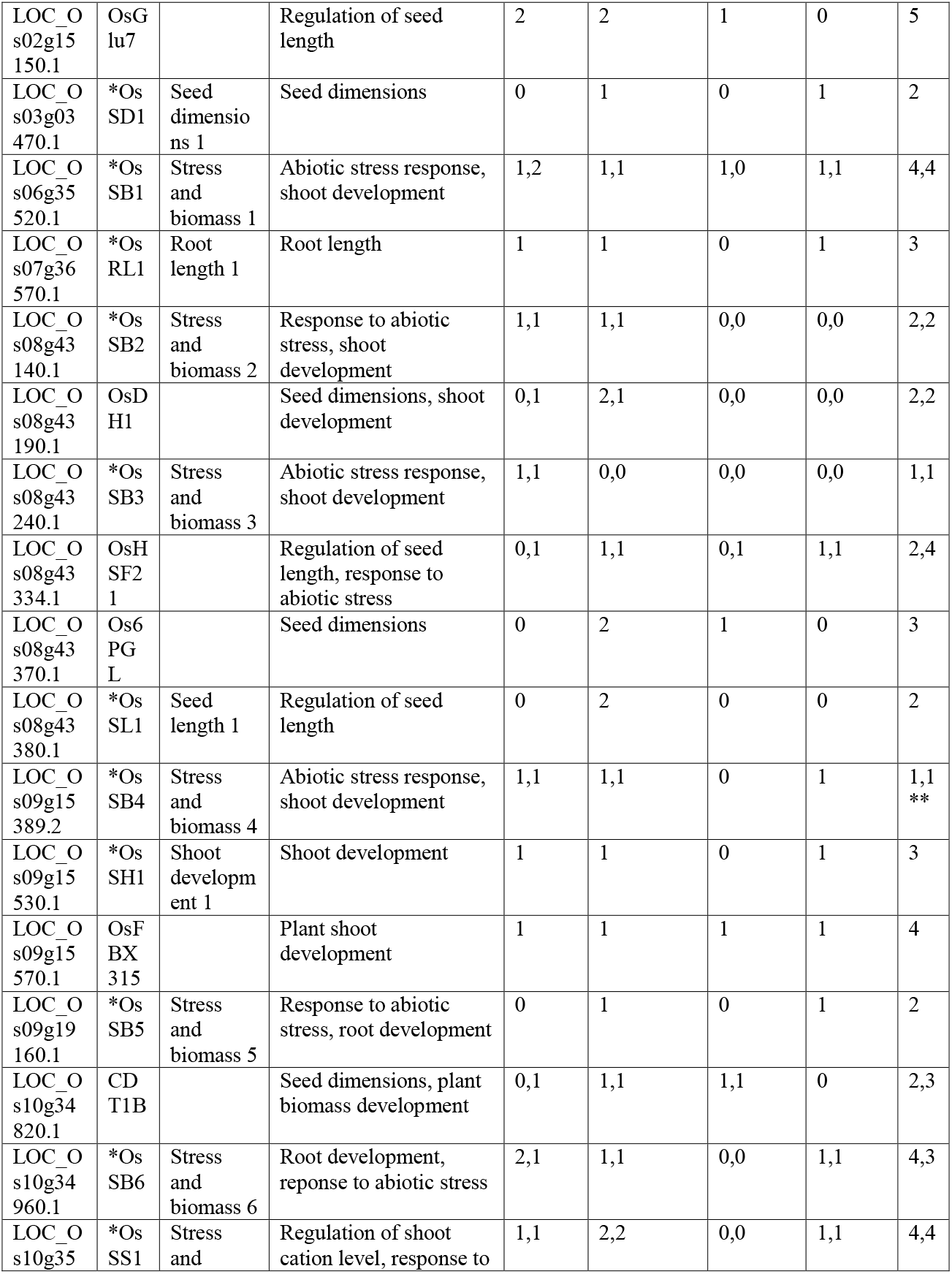
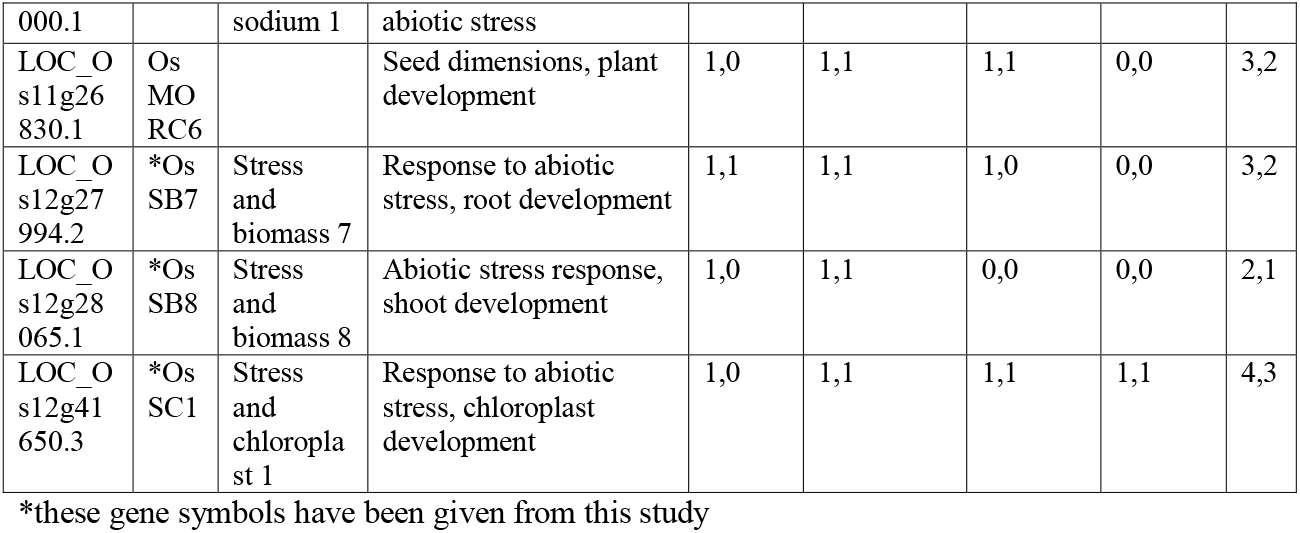
Elucidated gene roles.

## 4 Discussion

### 4.1 Informative features are hidden in observable phenotypes

It is evident that rice grains are not box-shaped objects. Nonetheless, our derived seed volume and seed density phenotypes found significant associations, some of which have been implicated in seed development in previous studies. Individual differences in plants of identical genotype are disturbingly disorderly. We found that variance within genotypes in our growth experiments were distinctly large for most traits other than seed properties. But such is customary to plant breeding experiments and many known and unknown factors are in play here. The question here is not how we can remove these confounding variables, since it is not only impossible but also contra-intuitive for practical purposes, but rather how we can derive informative features from observable phenotypes. To this end, researchers have devised complex phenotype models [50] or employed geometric methods such as multidimensional scaling to prepare testing phenotypes [51]. But more commonly, researches take ratios or differences and we have only employed these simple operations on top of our basic phenotypes. At the culmination of our study, our impression is that enumerating informative features from phenotypes for genomic studies in plants can be tricky and for best results, these experiments must be carefully planned. In light of this, we propose that highly heritable and better performing phenotypes can be subjected to complex modelling and geometric augmentation for extraction of more subtle information.

### 4.2 Biomass traits and chlorophyll content could be valuable indices for the screening of salinity tolerance

There lies significant ambiguity in the extrapolation of salt tolerance mechanisms in rice [52]. Gregorio et al. 1997 [18] popularized visual injury scoring for the screening of salinity tolerance. The application of SES for assessment of salinity tolerance has gathered some criticism since then [14]. We found the SES score to have meaningful correlations with all observed traits except for ion content (Supplementary figure 1). Our ion content traits are among the least heritable phenotypes (Table 1 and supplementary table 2) and their distributions deviate the most from a normal gaussian distribution (Supplementary figure 2). They are also the most variable across genotypes and among replicates (Supplementary table 3), least applicable for genomic prediction (Supplementary figure 3) and the least correlated between two test conditions and with the other phenotypes, including the composite SES score (Supplementary figure 1). There are however, significant near-binary differences in tissue sodium content for control and stress counterparts and the magnitude of this change greatly outweighs the variance observed across genotypes (Supplementary table 3). This proved that in spite of being the most significant physiological alteration with the most deleterious metabolic consequences, changes in sodium content for individual genotypes between control and stress conditions could not be an informative variable for prediction models. This lack of correlation and lack of reliable difference across genotypes was previously observed by Pires et al. 2015 [10]. Pires et al. also found that potassium content in roots and leaves were not significantly altered between control and stress group plants and thus, could not be a reliable predictor of salt tolerance in statistical models. There has already been ample criticism against the traditionalist application of the K^+^/Na^+^ ratio for evaluation of stress tolerance [14, 53] and our observations incur further scrutiny to the conventional practice of evaluating salinity tolerance across different genotypes using tissue ion content and K^+^/Na^+^ ratios. For these reasons, we deliberately did not derive any further phenotypes from sodium and potassium content traits.

Although ion content is linearly independent to salt injury, very high correlation between sodium and potassium content in roots under salt stress is observed (Supplementary figure 1). The lack of this correlation in shoots under stress conditions is an elegant demonstration of sodium exclusion from plant leaves for survival in salt stress. Chlorophyll A and B are not independent variables and although relatively less heritable, show high degree of correlation in both control and stress conditions and correlate strongly to salt injury. This was reported in Pires et al. 2015 and our findings confirm their observations. The relationship observed between biomass traits and salinity tolerance in our studies is highly noteworthy. It is always the combination of a number of salt tolerance strategies that a plant employs to cope with stress. Intuitively, salinity tolerance benchmarks are depicted by the ability of plants to accumulate and retain biomass and color. We observed that proportional losses of shoot and root biomass is instilled by salt stress. Whereas plant genotypes that had more chlorophyll in control conditions, did not retain more chlorophyll in stress conditions (Figure 3). Hence for chlorophyll, how much a plant has under normal conditions is not much relevant to how much a plant of the same genotype could maintain under salt stress conditions. How much biomass a plant of certain genotype retains in salt stress is however, linearly correlated to how much biomass it can put on in control conditions. Greater biomass in stress conditions correlates strongly and negatively to damage scores. It was also found by other researchers that the effect of salinity on biomass is more predominant in the shoots compared to roots [54]. With these observations, and on the basis of GWAS results, we place much emphasis on biomass traits and chlorophyll content for the assessment of salinity tolerance in rice.

### 4.3 GWAS results hint at reciprocity between chlorophyll and potassium ion traits

A large association block on chromosome 3 spanning in megabases was found significantly correlated with chlorophyll traits (git repository). Many loci for chlorophyll content are known to lie in the observed genomic range. The objective of our study was to identify genetic targets for molecular intervention. A lot of data had been generated and inclusion of the large block from chromosome 3 could have compromised the results by introducing many false positives. Likewise, from the GWAS studies, we emphasized genes and marker associations that had relevant annotations for traits showing significant associations. Some observed marker associations were empirically ignored to preserve the quality of the candidate batch, but they have all been included in supplementary table 8. Within the significant associations in supplementary table 8, three expressed transporter proteins belonging to the highly studied HAK family, HAK16, HAK21 and HAK27 were present. These genes have been implicated in salinity tolerance and are already considered as molecular targets for crop improvements. Markers within the transporter genes were associated with chlorophyll content and interestingly, some very important chlorophyll related genes were found to lie in the qCDP3 locus, which was discovered relating to shoot potassium content. An intricate mutual interdependence between shoot potassium and plant chlorophyll can be imagined which would explain why these signals did not reappear in stress condition where both chlorophyll levels and metabolically accessible potassium is compromised. However, this conjecture remains a farfetched hypothesis awaiting concrete scientific validation.

qCDP3 was associated with shoot potassium content in stress conditions but we categorized it as a stress tolerance related trait instead of a potassium content related trait. This was primarily done to simplify the interpretations of our findings, since throughout the experiment, a vast amount of data was generated and it presented a daunting task of manually curating and reviewing every significant hit. It did not escape our notice that multiple stress tolerance related traits, namely lost chlorophyll content and chlorophyll content in stress conditions were very close to the suggestive threshold for the same signal peak (refer to git repository for Manhattan plots). It is considered scientific malpractice to adjust significance thresholds to fit a promising narrative. The finding of a chloroplast localized potassium transporter and a set of compelling chloroplast related genes in qCDP3 would have been much more relevant in light of those associations that might exist, but did not cross the threshold barrier.

### 4.4 More on candidate genes in table 4

Final candidate selection was carried out under the following assumptions that, a: functionally impaired variants of important candidate genes exist in the population as copies in the genome and b: individuals carrying the genotype of the functional allele are quantifiably different in terms of phenotype. OsPRX86 is a peroxidase precursor protein belonging to the QTL qCDP7. The locus has been previously found to be associated to salt injury scores in rice [10]. A polymorphism causing a nonsense mutation in the gene was found to affect multiple salinity tolerance traits, namely, SES score, shoot length under stress and normal conditions and shoot weight. We characterize this protein as having an important role in salinity tolerance in rice. Another unannotated gene, LOC_Os09g15389.2, belonging to another previously identified locus [46], was observed to affect salt injury score, shoot weight and shoot length when the protein is truncated. Based on these observations, we impart a salinity tolerance role to this gene.

The SNP having the highest LD value in table 4 belongs to the QTL with the highest mean LD. OsPIL (MSU ID: LOC_Os12g41650.4) is a phytochrome interacting helix-loop-helix protein having 5 to 10-fold greater expression levels in plant leaves, and is known to play a part in abiotic stress response [55]. We suggest a role for this protein in the development of the photosynthetic apparatus in response to salt stress. Our observed SNP only affects transcript 4 out of 5 known transcripts. OsFBX315 (MSU ID: LOC_Os09g15570.1) is a lesser studied protein with no tagged ontology terms. We conclude that this protein plays an important part in plant shoot development and growth. We found a nonsense mutation in OsDH1 (MSU ID: LOC_Os08g43190.2) to significantly affect rice seed and shoot dimensions. OsDH1 is a metabolically active catalytic protein with up to 15-fold higher expression in seeds and embryos. The observed significant differences in both seed height and seed width, two seed traits that we found to share no significant correlation (supplementary figure 1), inspires us to suggest a novel role for the gene in seed dimension regulation. LOC_Os09g15530.1 is an unannotated gene which we found to affect shoot length and shoot weight in control condition as well as the visual SES score and shoot length in salt stress condition. We assume a functional role for this gene in rice shoot development and response to abiotic stress. We found OsHSF21 (MSU ID: LOC_Os08g43334) to affect grain length when impaired. OsHSF21 is known as a stress response gene [56], but our observation that the gene belongs to a cluster of genes having markedly higher expression in seeds and embryos makes us assume the involvement of this gene in early seed development. Two more genes, LOC_Os08g43380.1 and LOC_Os12g06160.1 are also lesser known and were found to affect seed length. LOC_Os03g03470.1 is another unannotated gene affecting seed length and seed height. We previously found seed length to have no correlation with seed height (Supplementary figure 1). He hypothesizes that this gene also governs seed dimension traits. LOC_Os12g28065 is an unannotated gene belonging to qCDP16, a QTL we could associate with shoot ion content. The effect on shoot length in stress conditions and the difference in shoot length between two conditions categorizes it as an abiotic stress response gene. LOC_Os09g19160 is an uncharacterized serine/threonine kinase, that belongs to a cluster housing another serine/threonine kinase gene, LOC_Os09g19160. These kinases were found to affect root length in normal and stress conditions. Similar kinases have been observed to play an important part in root development and nodulation [57]. We propose this role for these genes. Another lesser-known gene, LOC_Os08g43140.1 was found to influence shoot length in stress conditions. Glu7 (MSU ID: LOC_Os02g15150.2) is a glutelin gene known to influence seed morphology [58]. This gene also belongs to a locus found to be associated with seed traits. LOC_Os12g27994.2 is an uncharacterized decussate gene we found associated with root length in stress conditions that is known to play a part in phyllotaxy and plant organ development [59].

OsLTPL97 is an assumed protease inhibitor gene that was observed to affect shoot length in both normal and stress conditions. A mutation in the 6-phosphogluconolactonase gene, LOC_Os08g43370.1 disrupted seed width, seed length, seed height, root length and shoot weight in stress condition. We assign a ubiquitous and multifactorial role to this gene in plant biomass development. OsMORC6b (MSU ID: LOC_Os11g26830), is a Microrchidia protein involved in DNA repair and modification. An impairment of this protein was found to affect seed length, seed width, shoot length and root thickness. We conclude that this protein plays important regulatory functions in multiple plant tissues. CDT1B (MSU ID: LOC_Os10g34820) is a replication initiation protein. An impairment in CDT1B affects seed length, seed width and shoot length. LOC_Os10g34960.1 is an uncharacterized ubiquitin family protein. We delineate a functional role for the protein in root development.

## 5 Conclusion

Only by extrapolating the hidden diversity of forgotten landraces with the help of high-throughput molecular technologies will we be able to bioengineer the crops of the future that will thrive in substandard land without any compromise in yield. Equal emphasis on yield, tolerance and variability is essential for productive harvests and a food secure future. Quantifiable causal events on the genomic level are governed by countless external influences that affect polymorphisms on an evolutionary time scale. When we attempt to rapidly sieve these desirable attributes into our modern cultivars, it takes too long with breeding approaches and results are often far from expected with molecular approaches. A better understanding of the genomic landscape and the interplay between complex polygenic traits in plants is necessary for designing and implementing robust molecular protocols for phenotype augmentation. By analyzing multiple traits belonging to different categories in a heterogeneous plant population at once, we are able to assess the degree of heterogeneity in gene function and QTL marker effects. Common association signals for traits generally thought to be independent of one another demonstrates an interdependence among genetic factors. The findings of this study incite us to rethink our approaches in designing experiments for crop improvements. Narrow focus on one attribute at a time can only get us so far with phenotypes that are dictated by minute additive effects.

With rigorous GWAS analysis and stringent significance thresholds, we mapped 17 QTLs for various traits including 5 novel QTLs. We found reasonable evidence for novel gene roles in 21 candidate genes. Many of our trait and gene associations were found to overlap across intrinsically different phenotypes. Prior to GWAS, many traits sharing these very overlaps were observed to be linearly independent from one another. This leaves the nature of these causal overlaps to remain mostly unexplained, as they do not seem to be artifacts of population structure based on the comparable distributions in different subpopulations (Supplementary figure 2). These observations are however, not inconceivable. Deleterious changes in genes higher up in signaling cascades that occupy important central metabolic hubs will affect the functions of executor genes further down in the cascade. We affirm that we must be careful about the immediate interpretations of our findings as they have not yet been subjected to systematic case-control studies after gene cloning.

GWAS studies are convenient protocols for novel genetic discoveries. They have become even more popular with the subsistence and growth of publicly available genomic data. The automation of genomic studies is more relevant today because of recent advancements in automated mechanical and drone-based phenotype imaging technologies. It is our firm belief, that our experimental design which has been publicly documented on GitHub will aid future researchers and greatly facilitate GWAS and other studies in plant genomics.

## 6 Acknowledgements

The authors would like to wholeheartedly acknowledge Dr. Sabrina M. Elias from Independent University, Bangladesh and MU Sharif Shohan from the University of Dhaka for their valuable insights and guidance throughout the project. We acknowledge the Center for Bioinformatics Learning Advancement and Systematic Training (cBLAST) from the University of Dhaka and the University of Dhaka itself for enabling the necessary computational and experimental resources. We acknowledge Dr. Md. Sazzadur Rahman from Bangladesh Rice Research Institute (BRRI) for his assistance in multiplying seeds and preparing the germplasm. We also acknowledge the members of the Plant Biotechnology Lab at the University of Dhaka with special mention to Rabin Sarker and Raju Ahmed for help in handling plant material, screening for salinity tolerance and in enumerating and tabulating wetlab data. Finally, we acknowledge Anika Tahsin from the University of Dhaka for help in retrieving geolocational origins of the studied plant varieties.

## Notes

https://www.github.com/DeadlineWasYesterday/Cat-does-plant

### Competing Interest Statement

The authors have declared no competing interest.

## References

1. German Environment Agency, F.M.f.E.C.a.D., Ten million hectares of arable land worldwide are ‘lost’ every year: less and less fertile and healthy soil. 2015, Umweltbundesamt: Germany https://www.umweltbundesamt.de/en/press/pressinformation/ten-million-hectares-of-arable-land-worldwide-are.

2. Olesen, J.E., et al., Impacts and adaptation of European crop production systems to climate change. European Journal of Agronomy, 2011. 34(2): p. 96–112.

3. Gupta, G.S.J.R.E.S., Land degradation and challenges of food security. 2019. 11: p. 63.

4. Hossain, M.S.J.I.R.J.B.S., Present scenario of global salt affected soils, its management and importance of salinity research. 2019. 1: p. 1–3.

5. Eynard, A., R. Lal, and K. Wiebe, Crop Response in Salt-Affected Soils. Journal of Sustainable Agriculture, 2005. 27(1): p. 5–50.

6. Tsioumani, E., The State of the World’s Biodiversity for Food and Agriculture: a Call to Action? Environmental Policy and Law, 2019. 49: p. 110–112.

7. Solis, C.A., et al., Back to the Wild: On a Quest for Donors Toward Salinity Tolerant Rice. 2020. 11(323).

8. McCouch, S., et al., Feeding the future. 2013. 499(7456): p. 23–24.

9. Kromdijk, J. and S. Long, One crop breeding cycle from starvation? How engineering crop photosynthesis for rising CO 2 and temperature could be one important route to alleviation. Proceedings of the Royal Society B: Biological Sciences, 2016. 283: p. 20152578.

10. De Leon, T.B., S. Linscombe, and P.K. Subudhi, Molecular Dissection of Seedling Salinity Tolerance in Rice (Oryza sativa L.) Using a High-Density GBS-Based SNP Linkage Map. Rice, 2016. 9(1): p. 52.

11. Yu, J., et al., Genome-wide association study and gene set analysis for understanding candidate genes involved in salt tolerance at the rice seedling stage. Molecular Genetics and Genomics, 2017. 292(6): p. 1391–1403.

12. Zhao, Y., et al., New alleles for chlorophyll content and stay-green traits revealed by a genome wide association study in rice (Oryza sativa). Scientific Reports, 2019. 9(1): p. 2541.

13. Kawahara, Y., et al., Improvement of the Oryza sativa Nipponbare reference genome using next generation sequence and optical map data. Rice, 2013. 6(1): p. 4.

14. Pires, I.S., et al., Comprehensive phenotypic analysis of rice (Oryza sativa) response to salinity stress. 2015. 155(1): p. 43–54.

15. Mansueto, L., et al., Rice SNP-seek database update: new SNPs, indels, and queries. Nucleic acids research, 2017. 45(D1): p. D1075–D1081.

16. Amin, M., et al., Over-expression of a DEAD-box helicase, PDH45, confers both seedling and reproductive stage salinity tolerance to rice (Oryza sativa L.). Molecular Breeding, 2012. 30(1): p. 345–354.

17. Cock, J., S. Yoshida, and D.A. Forno, Laboratory manual for physiological studies of rice. 1976: Int. Rice Res. Inst.

18. Gregorio, G., D. Senadhira, and R. Mendoza, Screening rice for salinity tolerance, vol 22, IRRI discussion paper series. International Rice Research Institute, 1997.

19. IRRI, I.J.I.R.R.I., Philippine, Standard evaluation system for rice. 2002: p. 1–45.

20. Endelman, J.B., Ridge Regression and Other Kernels for Genomic Selection with R Package rrBLUP. 2011. 4(3): p. 250–255.

21. Fusi, N., et al., Warped linear mixed models for the genetic analysis of transformed phenotypes. Nature Communications, 2014. 5(1): p. 4890.

22. Browning, B.L., Y. Zhou, and S.R. Browning, A One-Penny Imputed Genome from Next-Generation Reference Panels. The American Journal of Human Genetics, 2018. 103(3): p. 338–348.

23. Raj, A., M. Stephens, and J.K. Pritchard, fastSTRUCTURE: Variational Inference of Population Structure in Large SNP Data Sets. 2014. 197(2): p. 573–589.

24. Bradbury, P.J., et al., TASSEL: software for association mapping of complex traits in diverse samples. Bioinformatics, 2007. 23(19): p. 2633–2635.

25. Letunic, I. and P. Bork, Interactive Tree Of Life (iTOL): an online tool for phylogenetic tree display and annotation. Bioinformatics, 2007. 23(1): p. 127–128.

26. Huang, M., et al., BLINK: a package for the next level of genome-wide association studies with both individuals and markers in the millions. GigaScience, 2019. 8(2).

27. Tang, Y., et al., APIT version 2: an enhanced integrated tool for genomic association and prediction. 2016. 9(2): p. 1–9.

28. Turner, S.D.J.B., qqman: an R package for visualizing GWAS results using QQ and manhattan plots. 2014: p. 005165.

29. Chang, C.C., et al., Second-generation PLINK: rising to the challenge of larger and richer datasets. GigaScience, 2015. 4(1).

30. Sakai, H., et al., Rice Annotation Project Database (RAP-DB): an integrative and interactive database for rice genomics. Plant Cell Physiol, 2013. 54(2): p. e6.

31. Trapnell, C., L. Pachter, and S.L. Salzberg, TopHat: discovering splice junctions with RNA-Seq. Bioinformatics, 2009. 25(9): p. 1105–11.

32. Trapnell, C., et al., Transcript assembly and quantification by RNA-Seq reveals unannotated transcripts and isoform switching during cell differentiation. Nat Biotechnol, 2010. 28(5): p. 511–5.

33. Zhang, C., et al., PopLDdecay: a fast and effective tool for linkage disequilibrium decay analysis based on variant call format files. Bioinformatics, 2019. 35(10): p. 1786–1788.

34. Lippert, C., et al., FaST linear mixed models for genome-wide association studies. Nature Methods, 2011. 8(10): p. 833–835.

35. Mai, N.T.P., et al., Discovery of new genetic determinants controlling the morphological plasticity in rice root and shoot under phosphate starvation using GWAS. 2020: p. 2020.10.31.363556.

36. To, H.T.M., et al., Unraveling the Genetic Elements Involved in Shoot and Root Growth Regulation by Jasmonate in Rice Using a Genome-Wide Association Study. Rice, 2019. 12(1): p. 69.

37. Zhang, J., et al., QTL mapping and candidate gene analysis of ferrous iron and zinc toxicity tolerance at seedling stage in rice by genome-wide association study. BMC Genomics, 2017. 18(1): p. 828.

38. Tao, Y., et al., Genome-wide association mapping of aluminum toxicity tolerance and fine mapping of a candidate gene for Nrat1 in rice. PLOS ONE, 2018. 13(6): p. e0198589.

39. Zhang, Y., et al., QTL identification for salt tolerance related traits at the seedling stage in indica rice using a multi-parent advanced generation intercross (MAGIC) population. Plant Growth Regulation, 2020. 92(2): p. 365–373.

40. Misra, G., et al., Whole genome sequencing-based association study to unravel genetic architecture of cooked grain width and length traits in rice. Scientific Reports, 2017. 7(1): p. 12478.

41. Huang, X., et al., Genome-wide association studies of 14 agronomic traits in rice landraces. Nature Genetics, 2010. 42(11): p. 961–967.

42. Cui, Y., F. Zhang, and Y. Zhou, The Application of Multi-Locus GWAS for the Detection of Salt-Tolerance Loci in Rice. 2018. 9(1464).

43. Frouin, J., et al., Tolerance to mild salinity stress in japonica rice: A genome-wide association mapping study highlights calcium signaling and metabolism genes. PLOS ONE, 2018. 13(1): p. e0190964.

44. Diop, B., et al., Bridging old and new: diversity and evaluation of high iron-associated stress response of rice cultivated in West Africa. Journal of Experimental Botany, 2020. 71(14): p. 4188–4200.

45. Dingkuhn, M., et al., Crop-model assisted phenomics and genome-wide association study for climate adaptation of indica rice. 2. Thermal stress and spikelet sterility. Journal of Experimental Botany, 2017. 68(15): p. 4389–4406.

46. Schläppi, M.R., et al., Assessment of Five Chilling Tolerance Traits and GWAS Mapping in Rice Using the USDA Mini-Core Collection. 2017. 8(957).

47. Rohila, J.S., et al., Identification of Superior Alleles for Seedling Stage Salt Tolerance in the USDA Rice Mini-Core Collection. 2019. 8(11): p. 472.

48. Delorean, E.E.J.-.-C.T. and Dissertations, Detecting durable resistance to rice bacterial blight. 2016.

49. Mather, K.A., et al., The extent of linkage disequilibrium in rice (Oryza sativa L.). 2007. 177(4): p. 2223–2232.

50. Al-Tamimi, N., et al., Salinity tolerance loci revealed in rice using high-throughput non-invasive phenotyping. Nature Communications, 2016. 7(1): p. 13342.

51. Yano, K., et al., GWAS with principal component analysis identifies a gene comprehensively controlling rice architecture. 2019. 116(42): p. 21262–21267.

52. Bhowmik, S., et al., Identification of salt tolerant rice cultivars via phenotypic and marker-assisted procedures. 2007. 10(24): p. 4449–4454.

53. GARCIA, A., et al., Sodium and potassium transport to the xylem are inherited independently in rice, and the mechanism of sodium: potassium selectivity differs between rice and wheat. 1997. 20(9): p. 1167–1174.

54. Negrão, S., et al., Recent updates on salinity stress in rice: from physiological to molecular responses. 2011. 30(4): p. 329–377.

55. Jeong, J., G.J.M. Choi, and cells, Phytochrome-interacting factors have both shared and distinct biological roles. 2013. 35(5): p. 371–380.

56. Muthuramalingam, P., et al., Global integrated omics expression analyses of abiotic stress signaling HSF transcription factor genes in Oryza sativa L.: An in silico approach. 2020. 112(1): p. 908–918.

57. Krusell, L., et al., Shoot control of root development and nodulation is mediated by a receptor-like kinase. Nature, 2002. 420(6914): p. 422–426.

58. Li, X. and T.W. Okita, Accumulation of Prolamines and Glutelins during Rice Seed Development: a Quantitative Evaluation. Plant and Cell Physiology, 1993. 34(3): p. 385–390.

59. Itoh, J.i., et al., Rice DECUSSATE controls phyllotaxy by affecting the cytokinin signaling pathway. 2012. 72(6): p. 869–881.

